# Structural basis for the RC–LH1 supercomplex of *Dinoroseobacter shibae* for anaerobic anoxygenic photosynthesis

**DOI:** 10.1101/2025.03.03.641346

**Authors:** Peng Wang, Ze-Kun Liu, Jian-Xun Li, Ying-Yue Zhang, Jing-Li Lv, Kang Li, Xiu-Lan Chen, Lu-Ning Liu, Yu-Zhong Zhang

## Abstract

Aerobic anoxygenic phototrophic (AAP) bacteria are essential for oceanic carbon cycling. However, the architecture and structural adaptations of their photosynthetic systems to ensure adequate light harvesting, electron transport, and oxidative resilience in oxygen-rich environments remain poorly understood. In this study, we present the 2.4-Å cryo-EM structure of the reaction center-light-harvesting 1 (RC–LH1) supercomplex from *Dinoroseobacter shibae* DFL-12, a marine AAP bacterial symbiont of benthic dinoflagellates. This RC–LH1 supercomplex features a closed LH1 ring comprising 17 αβ-subunits, each containing two spheroidenones per αβ-heterodimer—a previously unreported configuration in phototrophic bacteria. The cytochrome subunit of the RC is truncated to three hemes, in contrast to the four-heme configuration found in anaerobic relatives. The structure also reveals elongated bacteriochlorophyll (BChl) spacing, accounting for its blue-shifted absorption maximum that is optimized for low-light benthic environments. Furthermore, we identify a previously unknown protein-LRC (Light-Respiratory Connector), which structurally bridges the RC and LH1 components and exhibits homology to NADH oxidoreductase subunit E, suggesting a functional coupling between photochemical and respiratory electron transport. Collectively, these specific structural features allow AAP bacteria to balance anoxygenic photosynthesis and protection against oxidative damage, providing a mechanistic framework for them to thrive in oxygenated marine environments. Our study provides insights into the structural and functional variability of bacterial photosynthesis in response to oxygenated marine environments.

## Introduction

Photosynthesis, the biological process that converts solar energy into chemical energy, is fundamental to life on Earth and drives global biogeochemical cycles(1). Photosynthesis can be classified into oxygenic photosynthesis and anoxygenic photosynthesis. Oxygenic photosynthesis, primarily found in cyanobacteria and plants, releases oxygen through water splitting catalyzed by Photosystem II. In contrast, anoxygenic photosynthesis is distributed across phylogenetically ancient bacterial lineages (e.g., Proteobacteria, Chlorobi, Chloroflexi), which utilize alternative electron donors such as hydrogen sulfide, ferrous iron, or organic compounds for light-driven energy conversion, demonstrating metabolic versatility in adapting to anoxic environments(2). Among these organisms, aerobic anoxygenic phototrophic (AAP) bacteria occupy a unique ecological niche by performing anoxygenic photosynthesis in oxygen-rich environments(3). AAP bacteria decouple light harvesting from oxygen sensitivity, enabling photosynthesis exclusively under aerobic conditions while utilizing organic substrates for aerobic respiration(4).

AAP bacteria make up 1–10% of the total prokaryotes in the surface ocean, and play a crucial role in the ocean’s carbon cycling(3, 5, 6). This ecological prominence underscores their importance in the global carbon cycle and microbial food webs. Despite their wide distribution and significant role in marine ecosystems, the structural and mechanistic bases of their photosynthetic machinery remain poorly understood.

In anaerobic phototrophs including purple phototrophic bacteria (e.g., *Rhodobacter sphaeroides*) and green phototrophic bacteria (e.g., *Chlorobaculum tepidum*), the reaction center-light-harvesting 1 (RC–LH1) supercomplex functions as the central photosynthetic unit, orchestrating photon capture and cyclic electron transport. RC–LH1 supercomplexes exhibit remarkable architectural diversity across species(7). Except for the RC–LH1 containing double layered LH1 rings from *Gemmatimonas phototrophica*(8), RC–LH1 supercomplexes can be mainly divided into three categories: *Thermochromatium tepid*um(9–11), *Rhodospirillum rubrum*(12–14), *Blastochloris viridis*(15), *Rhodopila globiformis*(16), *Blastochloris tepida*(17), *Allochromatium tepidum*(18), *Roseospirillum parvum*(19) exhibits the RC encircled by a closed LH1 ring containing 17/16-subunit αβ-polypeptides, whereas *Rhodopseudomonas palustris*(20, 21), *Rba. sphaeroides*(22, 23), *Rba. blasticus*(24), *Rba. veldkampii*(25), and *Rba. capsulatus*(26) adopts a C-shaped LH1 array with an opening occupied by protein-W and PufX. Moreover, *Rba. sphaeroides*(23, 27, 28), *Rba. blasticus*(24), and *Rhodobaca bogoriensis*(29) can assemble RC–LH1 dimers with an S-shaped LH1 ring architecture. The structural diversity and plasticity of RC–LH1 underlies the evolutionarily conserved mechanisms that enable adaptation to different spectral ranges and the use of light intensity gradients across various light environments.

RC–LH1 structures are crucial for efficient energy and electron transfer in low-oxygen environments. However, AAP bacteria face a specific challenge: their photosynthetic apparatus operates in oxygenated environments where two distinct mechanisms generate reactive oxygen species (ROS). Triplet-excited bacteriochlorophylls (³BChls) produce ROS, whereas electron leakage from the photosynthetic electron transport chain to molecular oxygen exacerbates oxidative stress, collectively reducing photochemical efficiency(4).

Here, we report the cryogenic electron microscopy (cryo-EM) structure of the RC–LH1 supercomplex from *Dinoroseobacter shibae* DFL-12, a model marine AAP bacteria symbiont of benthic dinoflagellates. The structure reveals that the LH1 ring with elongated BChl distances is optimized for light capture in benthic habitats (short-wavelength light) with varying light intensities. A previously unidentified protein subunit (protein-LRC) is determined to bridge the RC and LH1, potentially integrating photosynthetic and respiratory electron transport. Moreover, the LH1 ring contains additional carotenoids and features a narrow quinone transfer channel, which could enhance ROS quenching and elevate the midpoint potential (E_m_) of the primary acceptor Q_A_ in the RC of AAP bacteria(4). Our study provides insights into aerobic anoxygenic photosynthesis and could inform the rational engineering of oxygen-tolerant photosynthetic systems.

## Results and discussions

### Overall structure of the RC–LH1 supercomplex

RC–LH1 supercomplexes were isolated from *D. shibae* cells cultivated phototrophically under aerobic conditions through sucrose density gradient ultracentrifugation (Fig. S1). Structural integrity of the purified complexes was confirmed by SDS-PAGE analysis (Fig. S1C). The cryo-EM structure of the *D. shibae* RC–LH1 supercomplex was resolved at 2.49 Å resolution (Fig. S2, Table S1), allowing clear visualization of amino acid side chains (Fig. 1A-C and S3) and enabling robust atomic model construction through iterative refinement (Fig. 2D-F).

**Fig. 1.**
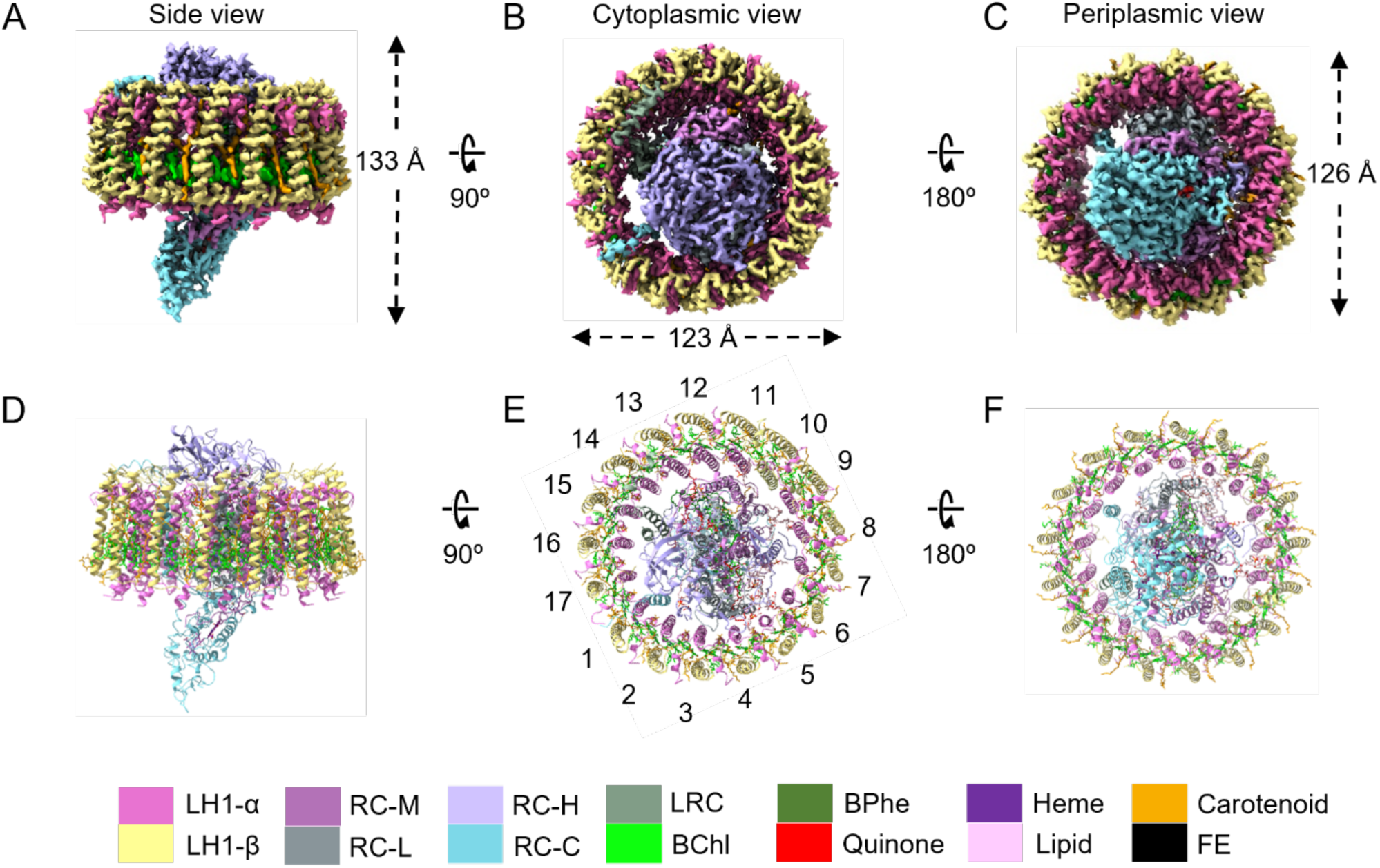
Cryo-EM structure of the RC-LH1 supercomplex from *D. shibae*. (A to C) Views of the RC–LH1 density map, colored as in the key at the bottom of the figure. (A) Side view of the RC–LH1 complex in the membrane plane, showing the height of the RC–LH1 complex. (B) Top view of the RC–LH1 supercomplex from the cytoplasmic side, showing the diameters of the long axes. (C) Bottom view of the RC–LH1 supercomplex from the periplasmic side, showing the diameters of the short axes. (D to F) Strutural model of the RC–LH1 complex in three different views corresponding to (A to C).

**Fig. 2.**
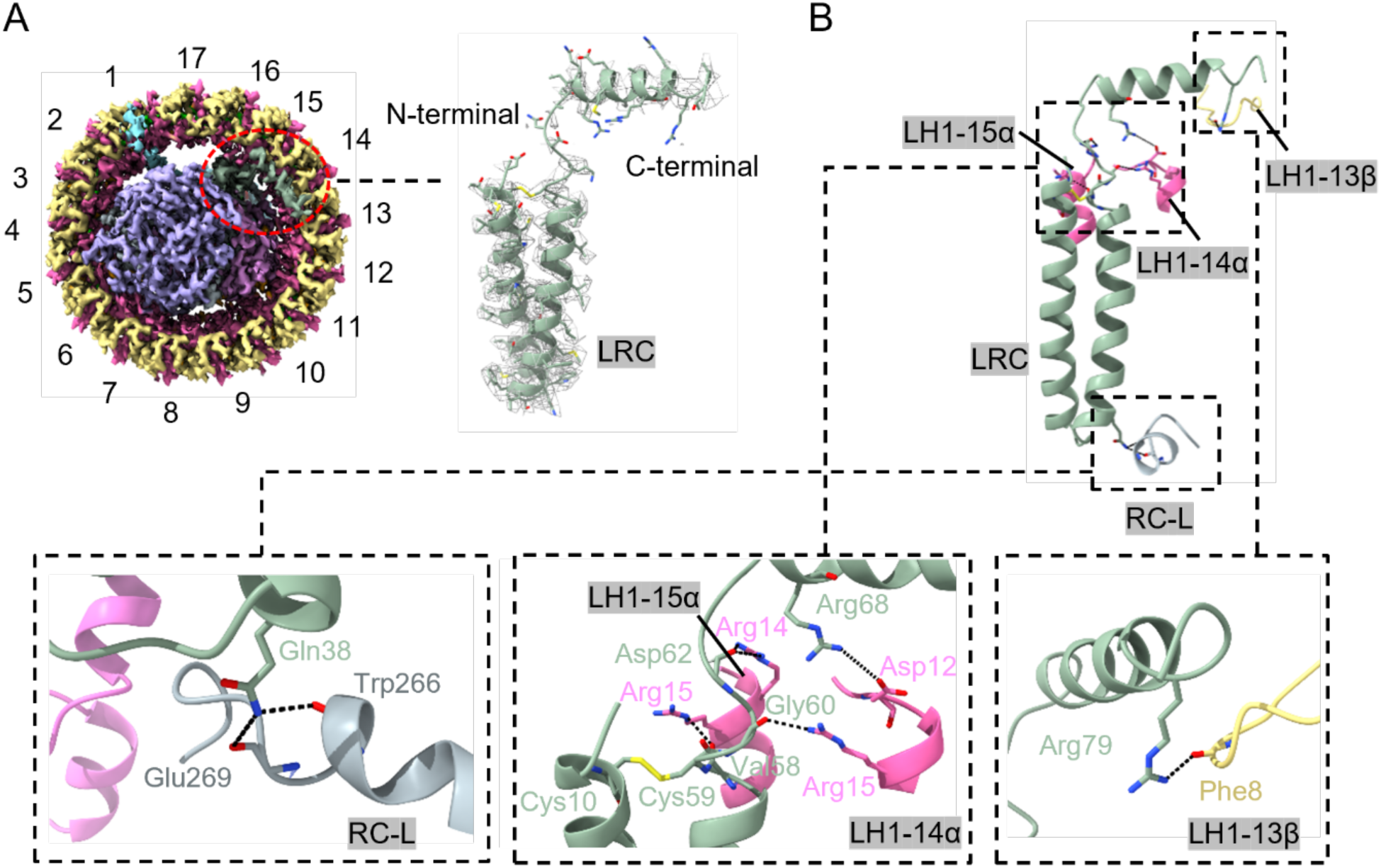
Structural analysis of protein-LRC. (A) The binding site and the electron density map of protein-LRC. (B) Interactions between protein-LRC and its surrounding subunits. Details are displayed by zoomed-in views.

The RC–LH1 supercomplex exhibits a slightly elliptical architecture in the projection view, with dimensions of 126 Å (short axis), 123 Å (long axis) (Fig. 1B-C), and 133 Å in vertical height (Fig. 1A). To our best knowledge, it represents the largest RC–LH1 structure with a single LH1 ring discovered to date. The RC core is encircled by a closed LH1 ring, which is composed of 17 αβ-heterodimers arranged circumferentially, each binding two bacteriochlorophyll *a* (BChl *a*) molecules and two spheroidenone carotenoids (34 total pigments per ring; Fig. 1D). The assignment of spheroidenones was corroborated by characteristic absorption spectra of the purified complexes (Fig. S1B) and previous biochemical studies(30). Structural modeling of the LH1 subunits revealed a conserved topology: both α- (51 residues) and β-polypeptides (44 residues) possess central transmembrane α-helices, with their N-terminal domains oriented toward the cytoplasmic membrane surface and C-termini extending periplasmically (Fig. 1D and S3).

The RC is composed of three core subunits (H, L, M) and a cytochrome c subunit that contains three hemes in contrast to the canonical four in anaerobic relatives (Fig. S3). Structural analysis identified five BChl *a* molecules, two bacteriopheophytin (BPhe *a*) molecules, one spheroidenone molecule, nine lipid molecules, three heme molecules, four ubiquinone-10 molecules, and a non-heme iron atom within the RC. Notably, we identified a previously unknown protein subunit, designated protein-LRC (Light-Respiratory Connector). The structural and functional implications of protein-LRC are discussed in the following sections.

The structure of the *D. shibae* RC–LH1 supercomplex resembles that of the *Blc. viridis* RC–LH1 supercomplex (PDB 6ET5)(15), both of which feature a closed LH1 ring with 17 αβ-polypeptides surrounding the RC. However, compared to the *Blc. viridis* counterpart, the *D. shibae* RC–LH1 supercomplex contains an additional carotenoid in each LH1 αβ-polypeptide, and features a larger LH1 ring diameter. More strikingly, *D. shibae* RC–LH1 contains a unique tri-heme cytochrome c subunit, a novel subunit protein-LRC, and lacks the γ-apoprotein present in *Blc. viridis* RC–LH1.

### The special protein-LRC

At a 2.49 Å resolution, the electron density map of the *D. shibae* RC–LH1 supercomplex revealed the presence of a previously unidentified protein subunit located between the RC and LH1 (Fig. 2A). However, no nucleotide sequences were found within the photosynthetic gene cluster, which encodes the protein that matches the electron density of this new subunit. The sequence of this specific protein was ultimately determined by matching electron density against the genomic protein sequence. This protein, designated protein-LRC, is hypothesized to be encoded by a gene within the NADH-quinone oxidoreductase gene cluster. In the RC–LH1 density map, only residues from Asp7 to Thr82 of the N-terminal domain of protein-LRC were resolved, adopting a “helix-turn-helix-turn-helix” fold (Fig. 2A). The two α-helices near the N-terminus are transmembrane helices, oriented parallel to the LH1 ring and situated between the RC and LH1-15αβ. The third α-helix extends along one side of the LH1 ring, from LH1-15αβ to LH1-13αβ. The predicted full-length structure of protein-LRC suggests that it consists of an N-terminal domain, a C-terminal domain, and a long, flexible connecting loop (Fig. S4). The lack of densities for the C-terminal domain and the connecting loop is likely due to their inherent flexibility or potential degradation/loss during sample preparation.

Protein-LRC forms extensive hydrogen bonds with neighboring LH1 α-subunits and the C-terminus of the RC-L subunit (Fig. 2B, Table S2). On the cytoplasmic side, heterodimeric bonds are formed between LRC-Asp62 and LH1-15 α-Arg14, LRC-Val58 and LH1-15 α-Arg15, LRC-Gly60 and LH1-13 α-Arg15, LRC-Arg68 and LH1-14 α-Asp12, as well as LRC-Arg79 and LH1-13 β-Phe8. On the periplasmic side, LRC-Gln38 forms hydrogen bonds with L-Trp266 and L-Glu269. Additionally, the residues LRC-Cys10 and LRC-Cys59 stabilize the spatial arrangement of the two transmembrane helices of protein-LRC by forming disulfide bonds.

Bioinformatic and structural analysis revealed that the C-terminus of protein-LRC exhibits high sequence and structural similarity to the C-terminal domain of Clade A NADH-quinone oxidoreductase subunit E (Fig. S5 and S6). In certain photosynthetic bacteria, both Clade A and Clade E NADH-quinone oxidoreductases coexist, whereas in others, only one type is present. The main distinction between them lies in the presence of an additional C-terminal domain of NADH-quinone oxidoreductase subunit E found in Clade A, which is not found in Clade E. These two NADH-quinone oxidoreductases are responsible for distinct adaptive responses to various environmental conditions(31). *Rba. sphaeroides* contains both types of NADH-quinone oxidoreductases and regulates their expression depending on growth conditions, whereas *Rba. capsulatus* only contains the Clade A type but can cleave the extra C-terminal domain of subunit E under specific growth conditions to adapt to its environment(32, 33). These findings suggest that the C-terminal domain of subunit E plays a crucial role in modulating the function of NADH-quinone oxidoreductase, implying that protein-LRC is likely involved in regulating NADH-quinone oxidoreductase activity.

Although the specific function of protein-LRC requires further investigation, the current findings allow us to hypothesize that protein-LRC likely serves as a link between the RC–LH1 and NADH-quinone oxidoreductase, potentially playing a crucial role in the coupling of photochemical and respiratory electron transport. Under anaerobic conditions, the high Q_A_ E_m_ in AAP bacteria(4) could hinder effective photosynthesis, whereas under aerobic conditions, this coupling may lead to the reduction of Q_A_ E_m_ in AAP bacteria, which promotes anoxygenic photosynthesis and cell growth.

### The special cytochrome c subunit

There are two types of RCs in photosynthetic bacteria: one features a tightly bound cytochrome c subunit on the periplasmic side that donates electrons to the photo-oxidized RC core, whereas the other lacks this cytochrome c subunit and accepts electrons directly from water-soluble carriers(34). Structural studies of RCs in several photosynthetic bacteria have shown that the cytochrome c subunit typically contains four c-type hemes aligned along its long axis(10, 15). In contrast, the cytochrome c subunit in *D. shibae* contains only three hemes—specifically, Hemes 2, 3, and 4, with Met and His residues serving as the sixth axial ligands for the heme irons—while the heme near the periplasmic side (heme 1) is absent (Fig. 3A and 3B). This observation is consistent with bioinformatic analysis, which reveals that the first heme-binding motif (Cys-Xaa-Xaa-Cys-His) is missing in the sequence of the cytochrome c subunit from *D. shibae* (Fig. 3A and S7).

**Fig. 3.**
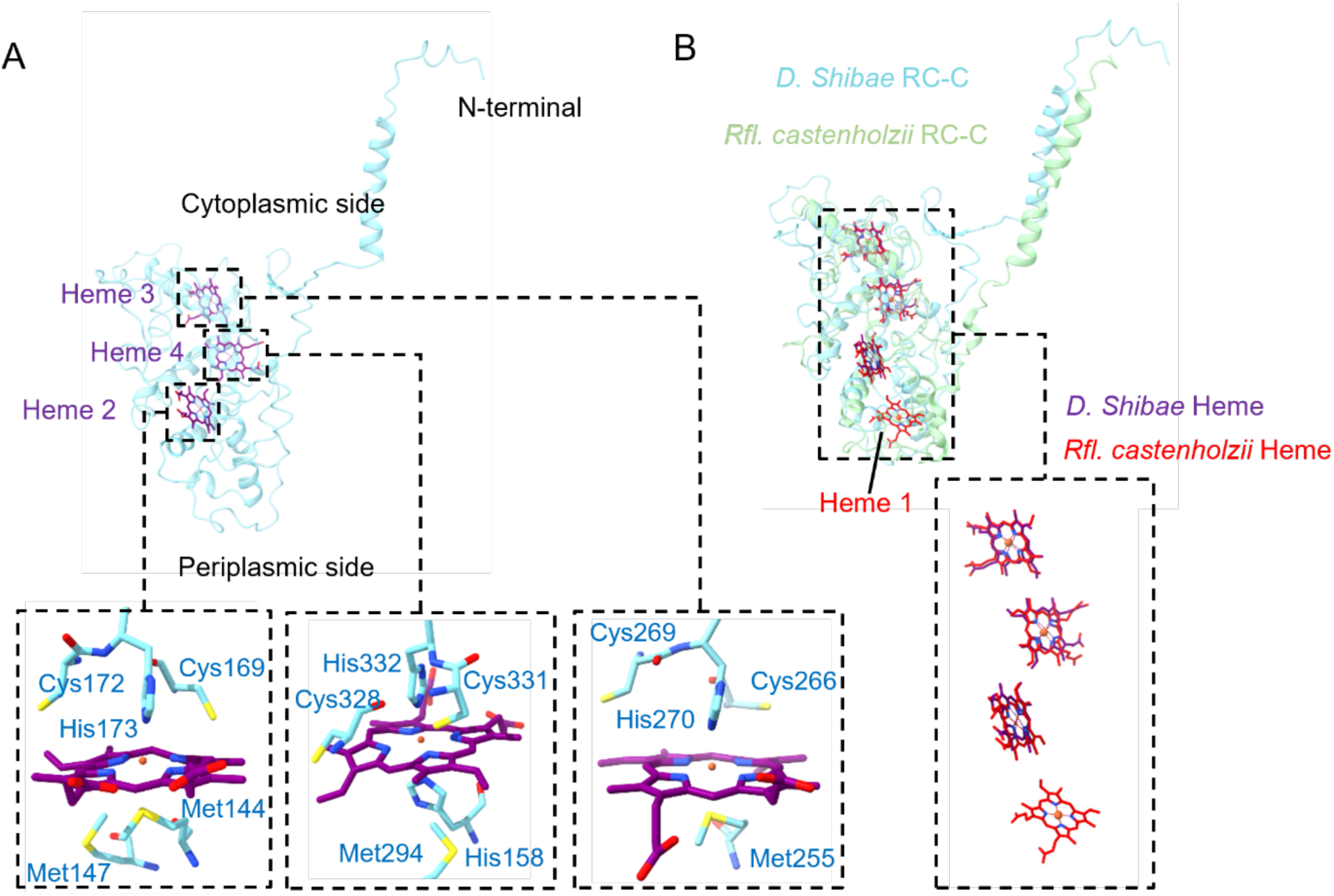
Structural analysis of the cytochrome c subunit. (A) The arrangement of the heme group in the cytochrome c subunit and its interactions with adjacent residues. Details are displayed by zoomed-in views. (B) Comparison between the cytochrome c subunits of *D. shibae* and *Rfl. castenholzii*. Details are displayed by zoomed-in views.

The loss of Heme 1 results in the absence of the traditional cytochrome *c*_2_ binding site. Therefore, the electron transfer pathway of this unique cytochrome c subunit should be remarkably different from that of the 4-heme cytochrome c subunit. Its electron transfer pathway and relationship with environmental adaptation still require further investigation.

In addition to the unique heme arrangement, the N-terminus of the cytochrome c subunit also exhibits distinctive features. It is attached to the LH1 ring and forms hydrogen bonds with LH1-1αβ. Heterodimeric bonds are formed between C-Lys12 and LH1-1 α-Arg14, as well as between C-Asp16 and LH1-1 α-Arg15 (Fig. 4C; Table S4). This creates an additional interaction beyond the typical RC– LH1 ring interactions, and likely provides a suitable target for the assembly of the RC–LH1 ring. The assembly of *D. shibae* RC–LH1 is further discussed below.

**Fig. 4.**
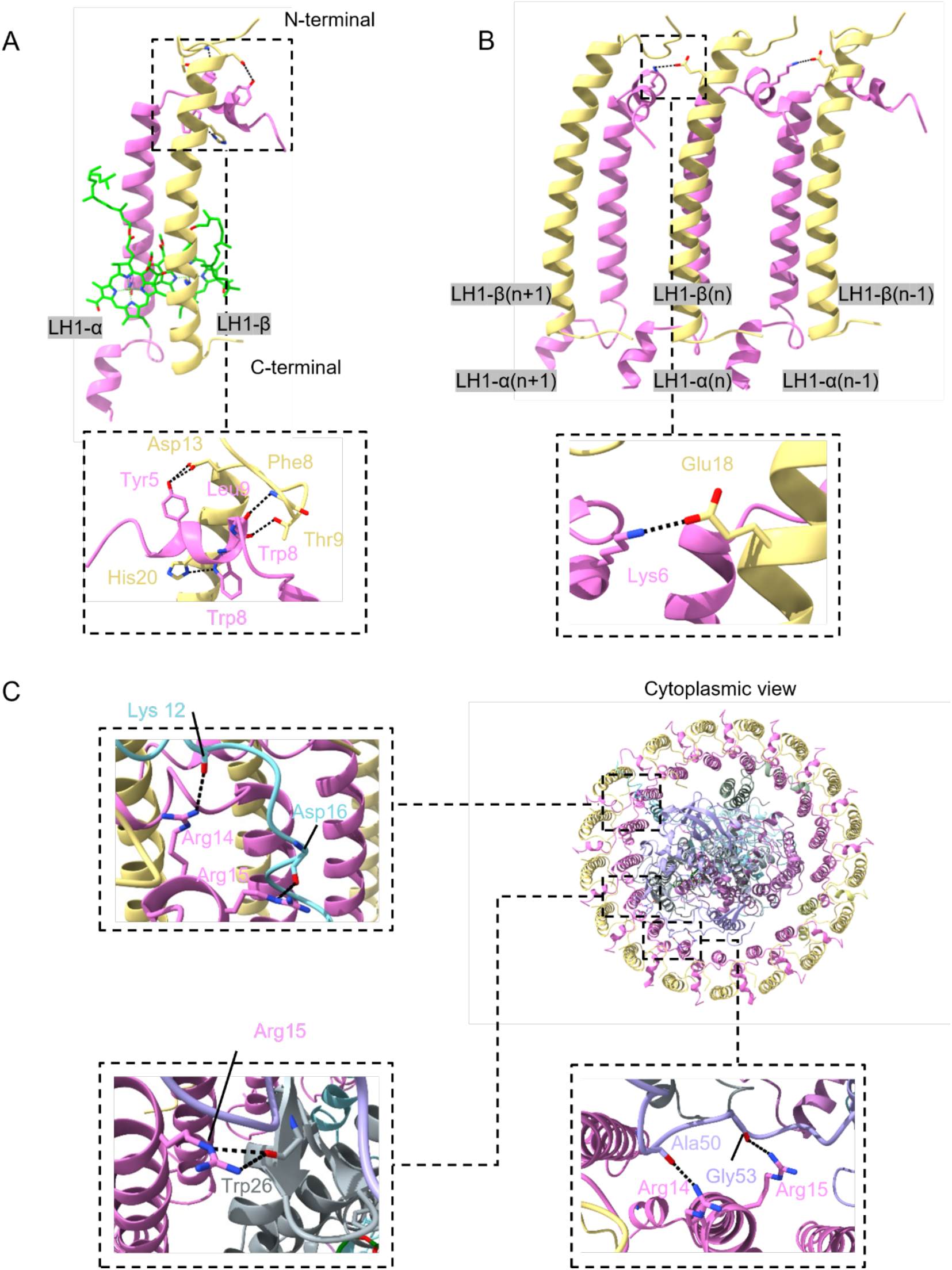
Interactions within the LH1 ring and between the RC and LH1 ring. (A) Hydrogen bond interactions within an LH1 subunit. (B) Intra- and inter-subunit hydrogen bonds within three LH1 αβ-subunits. (C) Hydrogen bond interactions between RC and LH1.

### Interactions within the LH1 ring and between RC and LH1

The interactions within the LH1 ring and between RC and LH1 of the *D. shibae* RC–LH1 supercomplex closely resemble those observed in other RC–LH1 supercomplexes. The structural integrity of the LH1 ring is maintained through an extensive hydrogen-bonding network, primarily localized within the N-terminal domains of the αβ-heterodimers (Figs. 4A and 4B; Table S3). Intra-heterodimer stabilization is facilitated by specific hydrogen bonds, including those between α-Tyr5 and β-Asp13, α-Leu9 and β-Phe8, as well as by interactions of α-Trp8 with both β-His20 and β-Thr9. Notably, the transmembrane α-helical regions of LH1 subunits do not participate in direct hydrogen bonding. Inter-heterodimer connectivity is established on the cytoplasmic face through hydrogen bonds linking α(n)-Lys6 of one heterodimer to β(n-1)-Glu18 of the adjacent subunit (Fig. 4B).

The interactions between RC and LH1 are mainly concentrated in three regions on the cytoplasmic side (Fig. 4C; Table S4). C-Asp16, L-Trp26, and H-Gly53 interact with Arg15 residues of neighboring LH1 α-subunits. In addition to these hydrogen bond positions, H-Ala50 and C-Lys12 interact with Arg14 residues of neighboring LH1 α-subunits. Together, these interfacial interactions anchor the RC core within the LH1 antenna complex while preserving structural flexibility that is essential for quinone transport.

### Assembly of RC–LH1

During our structural analysis of *D. shibae* RC–LH1, in addition to the density map for a complete RC–LH1 supercomplex, 3D classification also revealed two high-resolution density maps corresponding to *D. shibae* RC–LH1 supercomplexes with incomplete LH1 rings, termed RC–LH1-A and RC–LH1-B (Fig. 5A). Their structures were determined at resolutions of 2.44 Å and 2.75 Å, respectively (Fig. 5B and S2; Table S1).

**Fig. 5.**
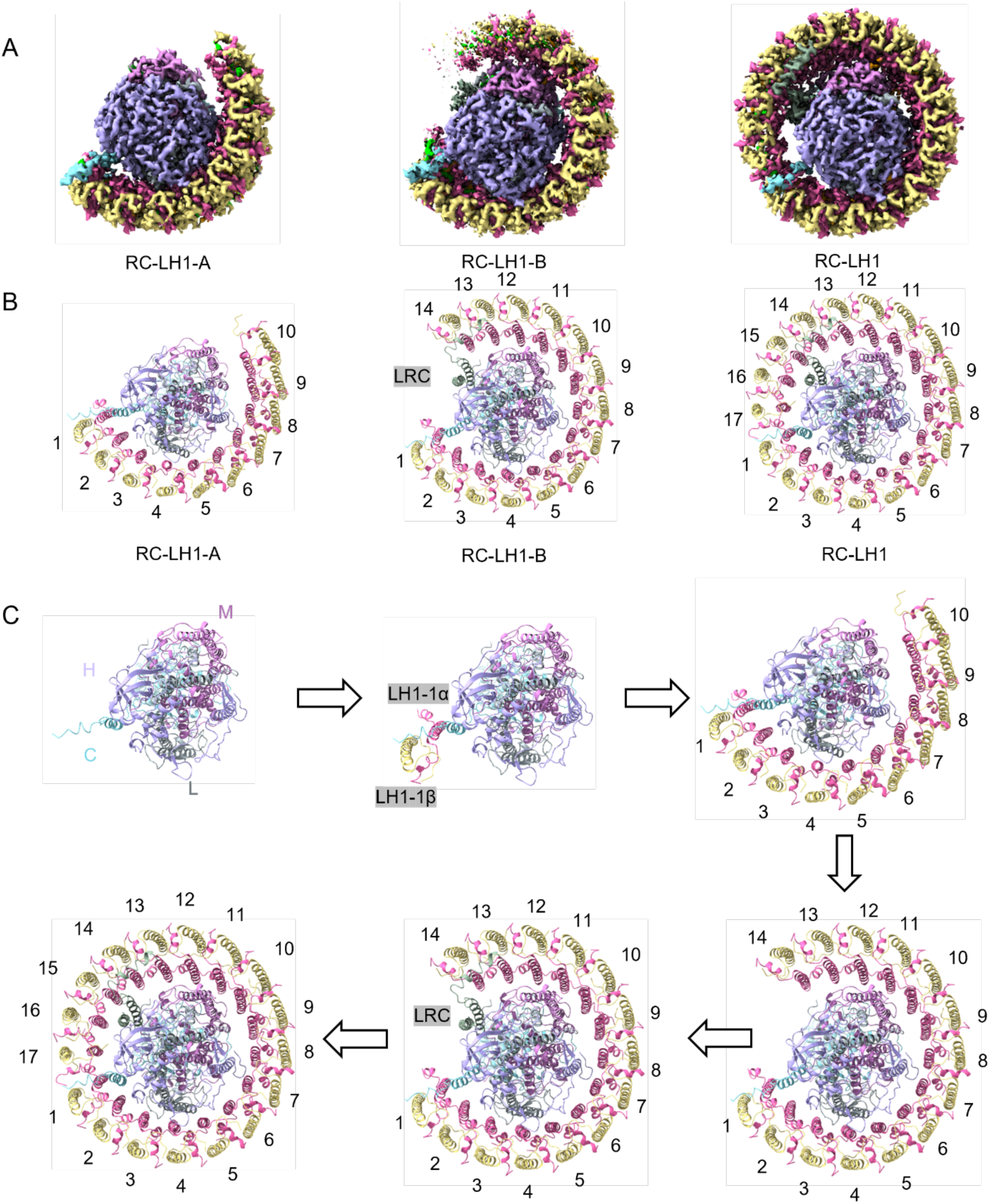
Schematic diagram of the assembly process of the *D. shibae* RC–LH1 supercomplex. (A) Density maps of RC–LH1 with complete and incomplete LH1 rings. (B) Structural models of RC– LH1 with complete and incomplete LH1 rings. (C) Schematic diagram of the assembly process of *D. shibae* RC–LH1 supercomplexes.

The RC structures in RC–LH1-A and RC–LH1-B are identical to those in the complete *D. shibae* RC–LH1. However, unlike the complete *D. shibae* RC–LH1 with 17 LH1 αβ-subunits, RC–LH1-A contains only 10 αβ-subunits, whereas RC–LH1-B has 14. Moreover, due to the absence of LH1-13 αβ and LH1-14 αβ, protein-LRC is also missing in RC–LH1-A, as the critical interactions required for its stabilization are no longer present.

The occurrence of RC–LH1-A and RC–LH1-B is likely due to the instability of αβ-subunit binding within the LH1 ring during the purification process. Interestingly, in both RC–LH1-A and RC–LH1-B, the LH1 ring always begins to lose subunits from LH1-17 αβ. This suggests that the LH1-1 αβ side has greater structural stability when interacting with the RC. As mentioned earlier, the N-terminal region of the cytochrome c subunit provides additional interactions between LH1-1 αβ and RC. This indicates that the N-terminal region of the cytochrome c subunit may function similarly to PufX in *Rba. sphaeroides*, serving as an anchoring site for the binding of the first LH1 αβ subunit(23).

These findings allow us to propose an assembly pathway for the *D. shibae* RC–LH1 supercomplex. Its assembly begins with the formation of the RC core, which is composed of the H, L, M, and cytochrome c subunits (Fig. 5C). Next, free LH1 αβ-subunits bind to the RC with the assistance of the N-terminal region of the cytochrome c subunit. The assembly of the LH1 complex is initiated by the incorporation of the first αβ-subunit, followed by the sequential binding of additional αβ-subunits around the RC, ultimately forming a closed ring. When the LH1 ring reaches the LH1-14 αβ position, protein-LRC begins to bind. (Figs. 5C). The precise mechanisms of this assembly and whether the proposed assembly process can be applied to many RC–LH1 supercomplexes similar to *D. shibae* RC–LH1. remain to be investigated.

### Arrangement of pigments and its adaption to aerobic habitats with varying light intensity

In the LH1 ring, the pigments form a tightly stacked array, systematically organized into two distinct groups: a ring consisting of 34 closely coupled BChls and another ring comprising 34 carotenoids (Fig. 6A). Due to the elliptical shape of the LH1 ring, the distance between BChls fluctuates, varying between 9.97 Å and 8.31 Å (Fig. S8). Both the Mg-Mg distances between BChls within individual LH1 αβ-heterodimers (9.65 Å on average) and between BChls of adjacent LH1 αβ-subunits (8.76 Å on average) (Fig. S8) are within 10 Å, ensuring efficient exciton coupling and energy resonance within LH1. We compared the BChl ring of the RC–LH1 supercomplexes of *D. shibae* and *Blc. viridis* (PDB 6ET5)(15), both of which contain 17 αβ-heterodimers. Our analysis showed that *D. shibae* RC–LH1 has a significantly larger radius and an increased average Mg-Mg distance (Fig. S9). The intra-subunit (8.8 Å) or inter-subunit (8.5 Å) Mg-Mg distances of BChl *b* in *Blc. viridis* RC–LH1 supercomplexare are relatively smaller, while *D. shibae* RC–LH1 has a larger average intra-subunit Mg-Mg distance of 9.65 Å, which exceeds those typically found in other RC–LH1 structures, such as those of *Tch. tepidum* (8.88 Å)(9), and *Rps. palustris* (9.53 Å)(20). The Mg-Mg distance within the BChl pairs reflects the degree of electronic coupling, with shorter distances indicating red shifts in the absorption spectrum and larger distances suggesting blue shifts(15). The enhanced Mg-Mg distance may account for the blue-shift in the absorption peak (866.5 nm) of BChls in LH1 from *D. shibae* (Fig. S1). Moreover, *D. shibae* RC–LH1 possesses BChl *a*, instead of BChl *b* that is present in *Blc. viridis* RC–LH1, and does not contain the γ-apoprotein that constrains the free movement of the LH1 ring and stabilizes the BChl *b* pairs(15). These structural variations are presumed to contribute to the distinct absorption spectra of the two RC–LH1 supercomplexes, with a blue shift in *D. shibae* and red shift in *Blc. viridis*(15).

**Fig. 6.**
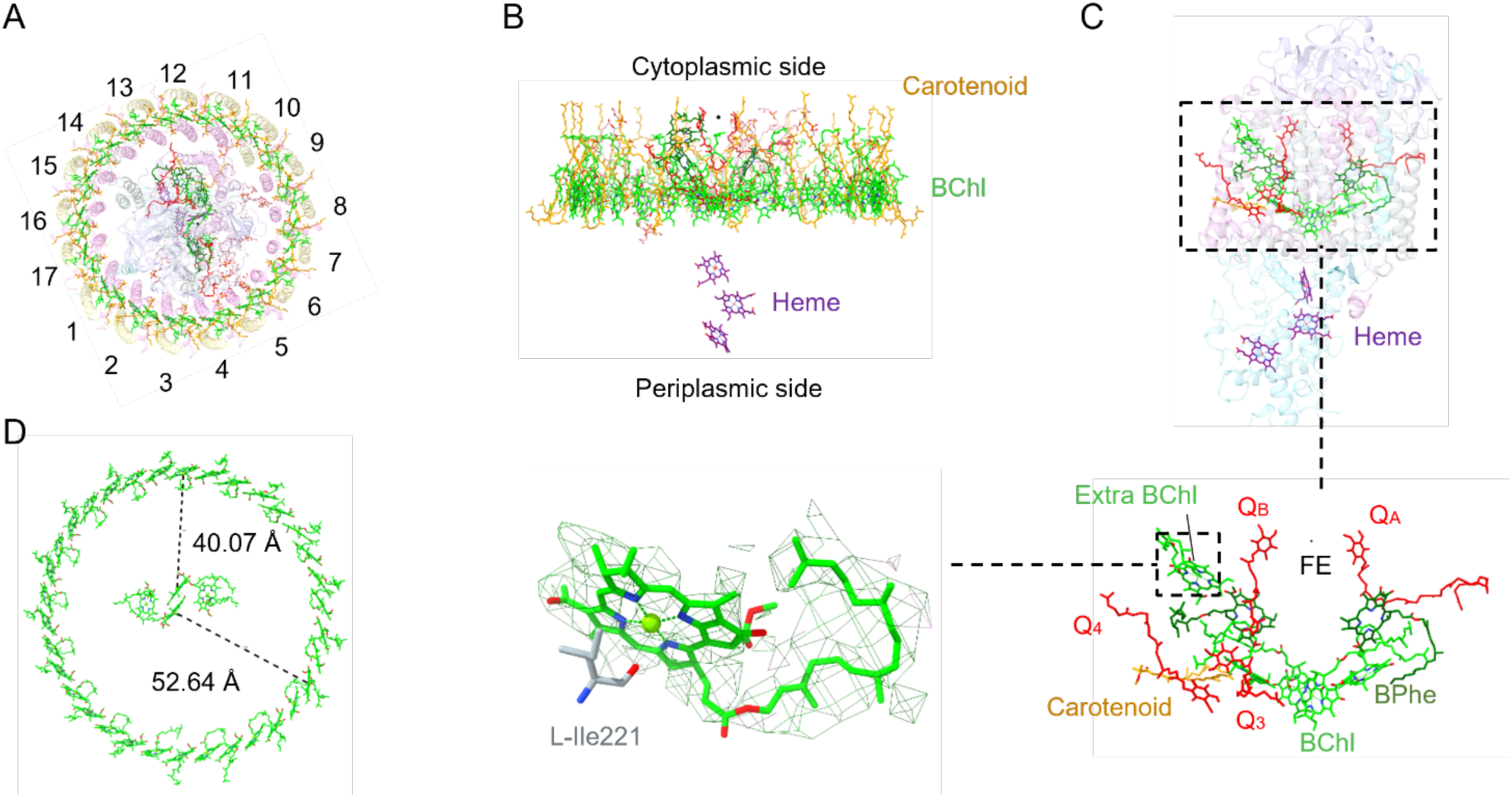
Arrangement of the pigments and cofactors in the *D. shibae* RC–LH1 supercomplex. (A) Pigments and cofactors viewed from the cytoplasmic side. (B) Side view of the arrangement of pigments and cofactors in the membrane plane. (C) Arrangement of pigments, quinones, and lipids associated with RC. (D) Maximum and minimum Mg-Mg distances between the BChls in the LH1 ring and the special pair BChls.

Moreover, the increased radius of the BChl ring results in an increase in the distances between the BChls in the LH1 ring and the special pair of BChls in the RC, ranging from 52.6 Å to 40.1 Å (Fig. 6D). The increased distance could reduce the efficiency of EET, potentially leading to the accumulation of ³BChl under high light conditions. This could result in the production and accumulation of ROS during aerobic photosynthesis. Compared to *Blc. viridis*, *D. shibae* thrives in oxygen-rich environments and features a larger LH1 ring with a greater diameter, making it more susceptible to severe oxidative stress(35). Consequently, specific protective mechanisms are necessary to mitigate oxidative damage. A notable structural difference is that each LH1 subunit in *D. shibae* RC–LH1 contains two carotenoid molecules, whereas *Blc. viridis* has only one. The primary carotenoid in *D. shibae* is spheroidenone, which is produced through the oxidation of spheroidene and OH-spheroidene via spheroidene monooxygenase (CrtA) under aerobic conditions. Typically, only one carotenoid is located between the αβ-polypeptides of the closed LH1 ring structure, facilitating the formation of channels for quinone trafficking across the LH1 αβ-polypeptides. However, *D. shibae* RC–LH1 incorporates two carotenoids per LH1 αβ-polypeptide within the closed LH1 ring. This adaptation enhances ROS quenching under high-light conditions and increases the number of light-harvesting pigments under low-light conditions, thereby improving light capture efficiency.

As we described above, the cofactors in the RC included five BChls *a*, two BPhes *a*, one spheroidenone, nine lipids, four ubiquinone-10 molecules, three hemes and one iron atom (Fig. 6C). The composition and arrangement of pigments in the *D. shibae* RC are consistent with those of most purple phototrophic bacteria, whereas the major differences lie in the presence of an additional BChl in the RC–LH1 supercomplex and the absence of a heme molecule. A similar additional BChl has also been observed in the RC–LH1 supercomplex of *Rsp. rubrum*(12, 13), but there is a significant difference in its positioning within the RC (Fig. 6C). The special BChl in *D. shibae* RC–LH1 is located between the RC-H, RC-L subunits, and LH1-1αβ, where it stabilizes the spatial structure by interacting with the main chain oxygen of L-Ile221 and is positioned closer to the BPhe *a* on the non-electron-transmitting side (Fig. 6C). The extra BChl may play a role in enhancing the efficiency of electron transfer (EET) from the LH1 ring to the RC, which increases EET efficiency under low light. Moreover, under high-light conditions, this additional BChl may dissipate excess energy through non-radiative decay, preventing photodamage and acting as an electron reservoir to store electrons transferred from the active center, thus alleviating the generation of reactive ROS.

### Pathways of quinone/quinol exchange

The special pair of BChls acts as the primary electron donor and is excited to P880*; this excited state then transfers an electron to BChl *a* and BPhe *a*, and subsequently to Q_A_. Q_B_ accepts electrons from Q_A_. Once quinone receives two electrons, it binds two protons from the cytoplasm and escapes from the RC to the neighboring Cyt *bc*_1_. In RC–LH1s with open LH rings, quinones/quinols are assumed to primarily shuttle through the gap in the LH1 ring, such as *Rba. sphaeroides*(23), *Rba. veldkampii*(25), and *Rba. capsulatus*(26). In contrast, in RC–LH1s with closed LH rings forming a palisade around the RC, quinones/quinols are proposed to diffuse through relatively small channels between two adjacent LH1 αβ subunits.

Four UQ-10 molecules were identified in the density map of *D. shibae* RC–LH1 (Fig. 6C and 7A). Two UQ-10 molecules function as the primary (Q_A_) and secondary (Q_B_) quinone acceptors (Fig. 7A). The head of Q_A_ is hydrogen bonded to His220 and Ala261 residues of RC-M, and the head of Q_B_ is hydrogen bonded to RC-L residues His191, Ser224, Ile225, and Gly226 (Fig. S10; Table S5). Two additional putative UQ-10 molecules (Q_3_ and Q_4_) are located in the gap between LH1 and the RC and are mainly surrounded by nonpolar residues of RC-L, RC-M, and protein-LRC. Q_3_ shares the same orientation as Q_B_ and is in close proximity to the isoprenoid tail of Q_B_ (Fig. 7A), suggesting that Q_3_ is in an appropriate position for the exchange of Q_B_ after double reduction and protonation.

**Fig. 7.**
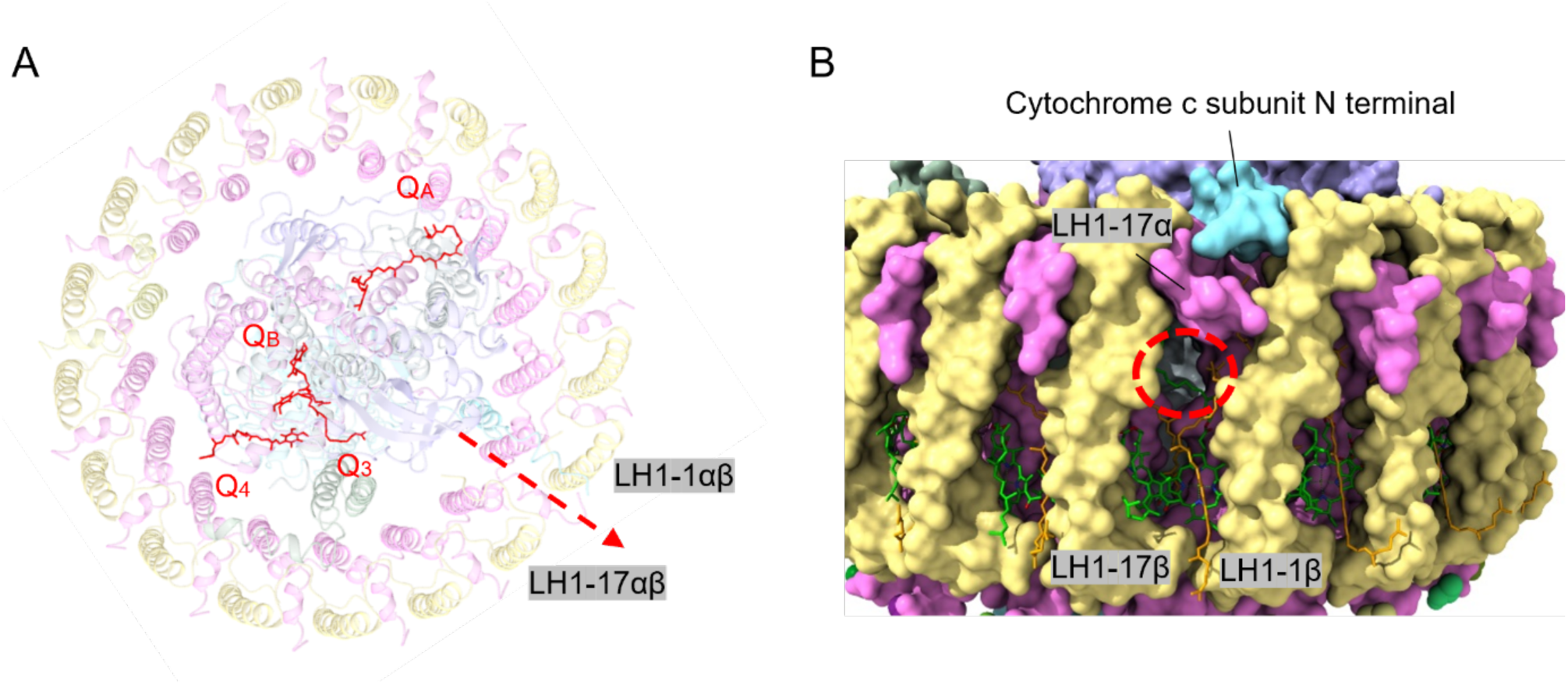
Proposed quinone/quinol diffusion channel in the *D. shibae* RC–LH1 supercomplex. (A) Cytoplasmic view. (B) Side view. Possible channels in the LH1 ring are indicated by red dashed circles.

Owing to the presence of additional carotenoid molecules in the LH1 ring, the gaps between most LH1 αβ subunits appear too small to accommodate quinone passage. Only a narrow pore exists between LH1-1αβ and LH1-17αβ, barely allowing quinone molecules to pass through (Fig. 7B). This pore formation is facilitated by the binding of the N-terminal domain of the cytochrome c subunit, which induces conformational changes in the N-terminal regions of LH1-17α and effects the binding conformation of BChl *a* (Fig. S11 and S12). This pore is notably small (10.2 Å in diameter) compared to that in *Blc. viridis* RC–LH1 (13.4 Å in diameter) (Fig. S13). This may be related to a slower electron transfer and a higher E_m_ of Q_A_ in *D. shibae* like AAP bacteria.

### Conclusions

The cryo-EM structure of the *D. shibae* RC–LH1 supercomplex unveiled key structural innovations driving its adaptation to aerobic anoxygenic photosynthesis. The cytochrome subunit of *D. shibae* RC–LH1 is integrated with three hemes instead of four hemes as found in anaerobic relatives. The LH1 ring exhibits elongated BChl spacing, a configuration likely optimized for efficient light harvesting in the fluctuating light conditions of benthic ecosystems. The tightly closed LH1 architecture is enriched with carotenoids and features a constricted quinone channel. More strikingly, a previously unidentified bridging subunit (protein-LRC) physically connects the RC and LH1, offering a potential molecular conduit for coordinating photosynthetic electron transport with respiratory pathways. These structural features provide the mechanistic basis of oxygen-tolerant photosynthesis in AAP bacteria and highlight their physiological adaptations in oxygen-dominated environments.

## Materials and Methods

### Growth condition of *D. shibae*

Wild-type *Dinoroseobacter shibae* DFL-12 (DSM-16493) was obtained from DSMZ. *D. shibae* cells were grown phototrophically under aerobic conditions in liquid 2216e medium at 30°C in glass bottles under a light intensity of 25 μmol photons s^−1^m^−2^ (Bellight 70 W halogen bulbs).

### Purification of RC−LH1 complexes

Cells were harvested by centrifugation at 5,000 × *g* for 10 min at 4°C, washed three times with Tris-HCl (pH 8.0), and resuspended in 20 mM HEPES (pH 8.0). The cells were disrupted by passage through a French press three times at 16,000 psi. Cell debris was removed by centrifugation at 20,000 × *g* for 30 min. Membranes were collected by centrifuging the resulting supernatant at 125,000 × *g* for 90 min and solubilized by the addition of β-DDM (n-dodecyl β-D-maltoside) to a final concentration of 3% (w/v) for 30 min to 60 min in the dark at 4°C with gentle stirring. Unsolubilized proteins were removed by centrifugation at 21,000 × *g* for 30 min. The supernatant was then applied onto 10–25% (w/v) continuous sucrose gradients made with working buffer containing 0.01% (w/v) β-DDM. Gradients were centrifuged at 230,000 × *g* for 18 h. The RC–LH1 complexes in the sucrose gradient solution were collected, and the purity of RC–LH1 complexes was characterized by sodium dodecyl sulfate-polyacrylamide gel electrophoresis (SDS-PAGE) and absorption spectra (Fig. S1).

### Absorption spectra

Purified RC–LH1 complexes were collected from sucrose gradients, and absorbance was measured from 300 to 900 nm at 1-nm intervals at room temperature using a Libra S22 spectrophotometer (Biochrom, United Kingdom).

### Cryo-EM data collection

A 4 μL aliquot of RC–LH1 sample was applied to a freshly glow-discharged holey carbon grid (Quantifoil Au R2/1, 200 mesh) with a continuous carbon support. The grid was blotted for 2 s at 100% humidity and 10°C, with a force level of 0, and immediately plunge-frozen into liquid ethane cooled by liquid nitrogen using a Vitrobot Mark IV (Thermo Fisher, USA). The grids were then loaded into a 300 kV Titan Krios G3i microscope (Thermo Fisher, USA) equipped with a Falcon 4 direct electron detector (Gatan, USA) for data acquisition. A total of 5962 movie stacks were automatically recorded using EPU (Thermo Fisher, USA)(36) at a total dose of 40 e^−^Å^−2^ per stack, with a defocus range of −0.8 to −1.8 μm and a pixel size of 0.97 Å.

### Data processing

Data processing was conducted using cryoSPARC (v4.4.1)(37). Patch motion correction and contrast transfer function (CTF) correction were performed, and micrographs with a resolution worse than 6 Å were discarded. From the remaining 5854 micrographs, 1,011,021 particles were picked using cryoSPARC’s Blob picking algorithm. The particles were extracted and subjected to three rounds of 2D classification, yielding 271,106 particles for further analysis. These particles were used for Ab initio reconstruction and 3D classification into four classes. Three ‘best-looking’ classes were selected for non-uniform refinement in cryoSPARC (v4.4.1), using the ‘optimize per-particle defocus’ option, yielded final maps with resolutions of 2.49 Å for the *D. shibae* RC–LH1 supercomplex, 2.44 Å for the RC–LH1-A supercomplex, and 2.75 Å for the RC–LH1-B supercomplex. Resolution estimates were based on the gold-standard fourier shell correlation at 0.143.

### Model building and refinement

The structure of the RC–LH1 supercomplex of *Blc. viridis* (DSM-133) (PDB ID: 6ET5) was initially docked into the cryo-EM map of the RC–LH1 complex using UCSF Chimera (v1.17)(38). The model was manually revised and refined based on cryo-EM density using Coot (v 0.9.4)(39), followed by real-space refinement using Phenix (v1.20.1)(40). Figures were generated using UCSF Chimera (v1.17) and ChimeraX (v1.16)(38, 40–42).

### Data and materials availability

The cryo-EM density map of *D. shibae* RC–LH1 supercomplex, RC–LH1-A supercomplex and RC– LH1-B supercomplex were deposited in the Electron Microscopy Data Bank (EMDB, www.ebi.ac.uk/pdbe/emdb/) under the accession code EMDB-63379, EMDB-63381 and EMDB-63382. Atomic coordinates of *D.* s*hibae* RC–LH1supercomplex, RC–LH1-A supercomplex and RC– LH1-B supercomplex were deposited in the Protein Data Bank (PDB, www.rcsb.org) with the accession code 9LTS, 9LTU and 9LTV. All data are available in the main text or supplementary material.

## Acknowledgments

We are grateful to Qiu-Yao Jiang (Shandong First Medical University, China) for cryo-EM data collection. We thank Xiao-Ju Li (Shandong University, China) for their assistance in TEM. This work was supported by the National Key R&D Program of China (2023YFA0914600, 2021YFA0909600, 2024YFC2816000), the National Natural Science Foundation of China (32330001, 32070109, 32170127, 32370136), Program of Shandong for Taishan Scholars (tspd20240806, tsqn202408064), the SKLMT Frontiers and Challenges Project (SKLMTFCP-2023-06).

## Author contributions

Y.-Z.Z. and L.-N.L. conceived the study. P.W., and L.-N.L. designed the experiments. P.W., Z.-K.L., Y.-Y.Z., J.-X.L., K.L., J.-L.L., M.-Q.W. performed the experiments. Z.-K.L., P.W., Y.-Y.Z. purified the protein samples and conducted optical spectral analysis. P.W., J.-X.L., K.L. collected cryo-EM data and processed the cryo-EM data. P.W. and Z.-K.L. assembled and generated the structural model and performed structural analysis. P.W., Z.-K.L., X.-L.C., Y.-Z.Z. and L.-N.L. wrote the manuscript. All the authors contributed to the discussion and improvement of the manuscript.

## Supplementary Information

**Fig. S1.**
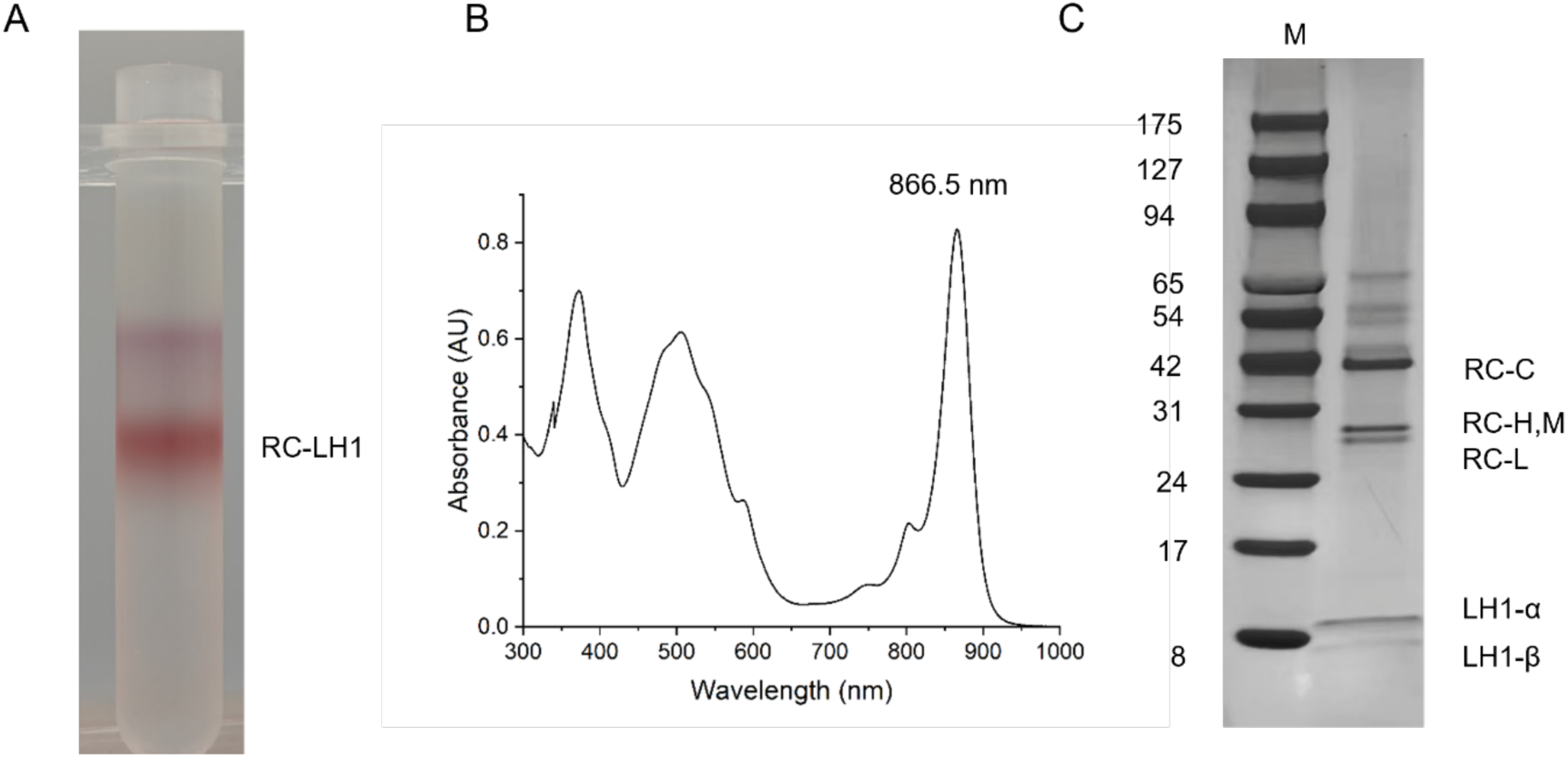
Purification and characterization of *D. shibae* RC–LH1 supercomplex. (A) Sucrose gradient ultracentrifugation result of photosynthetic membrane complexes from *D. shibae*. (B) Room-temperature UV-vis absorbance spectra of the purified sample (C) SDS-PAGE of purified sample.

**Fig. S2.**
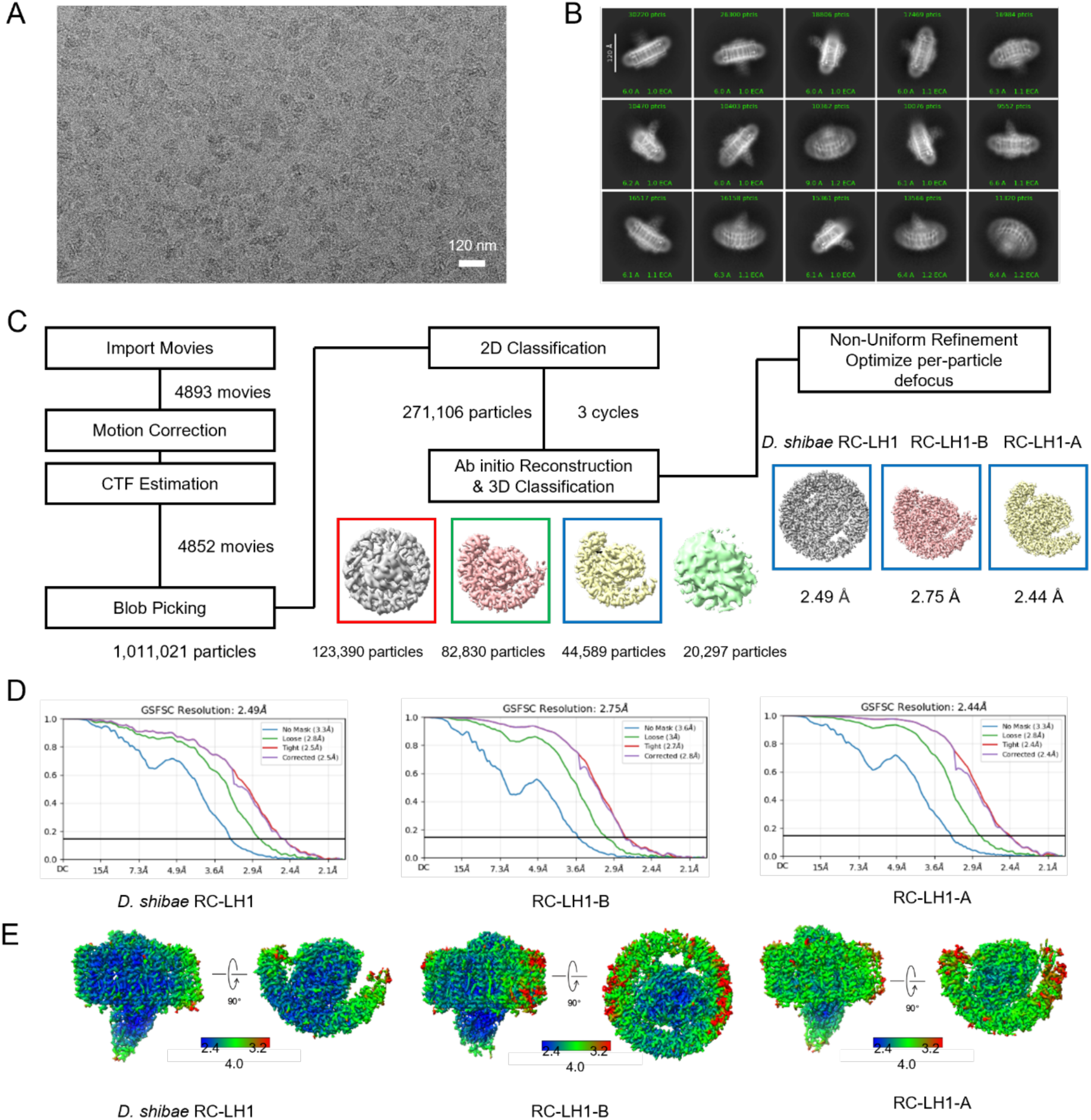
Cryo-EM data process of *D. shibae* RC–LH1 supercomplex. (a) Motion-corrected example of a cryo-EM captured movie. (b) Representative reference-free 2D class averages. (c) Overview of cryo-EM data processing. Selected 3D class that went into further processing are marked with rectangles. (d) Fourier Shell Correlation (FSC) curves generated by cryoSPARC. Global resolution values were calculated according to the gold-standard FSC = 0.143. (e) Local resolution of the cryo-EM map.

**Fig. S3.**
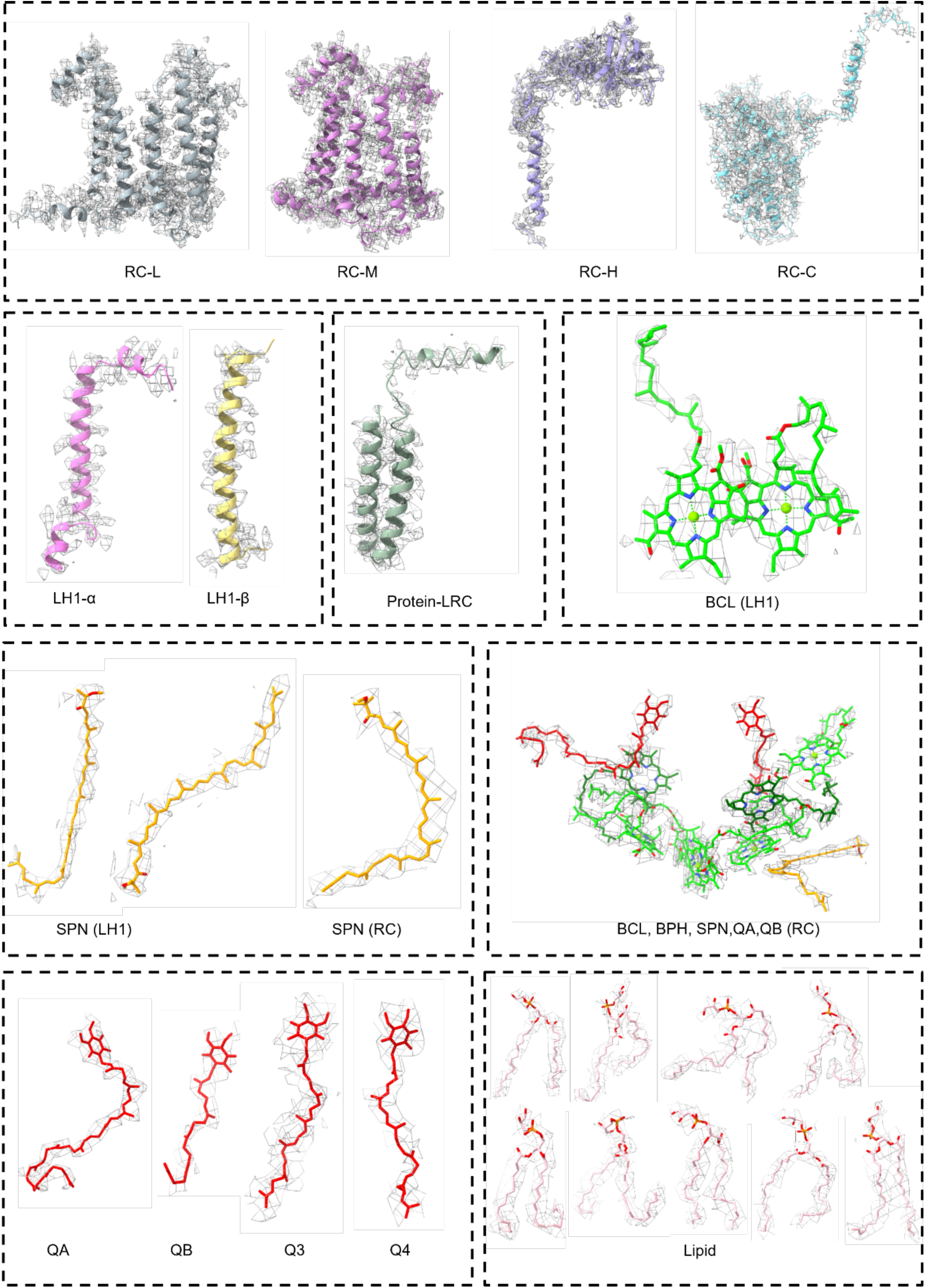
Cryo-EM map densities and structural models of protein peptides and cofactors in the *D. shibae* RC–LH1 supercomplex.

**Fig. S4.**
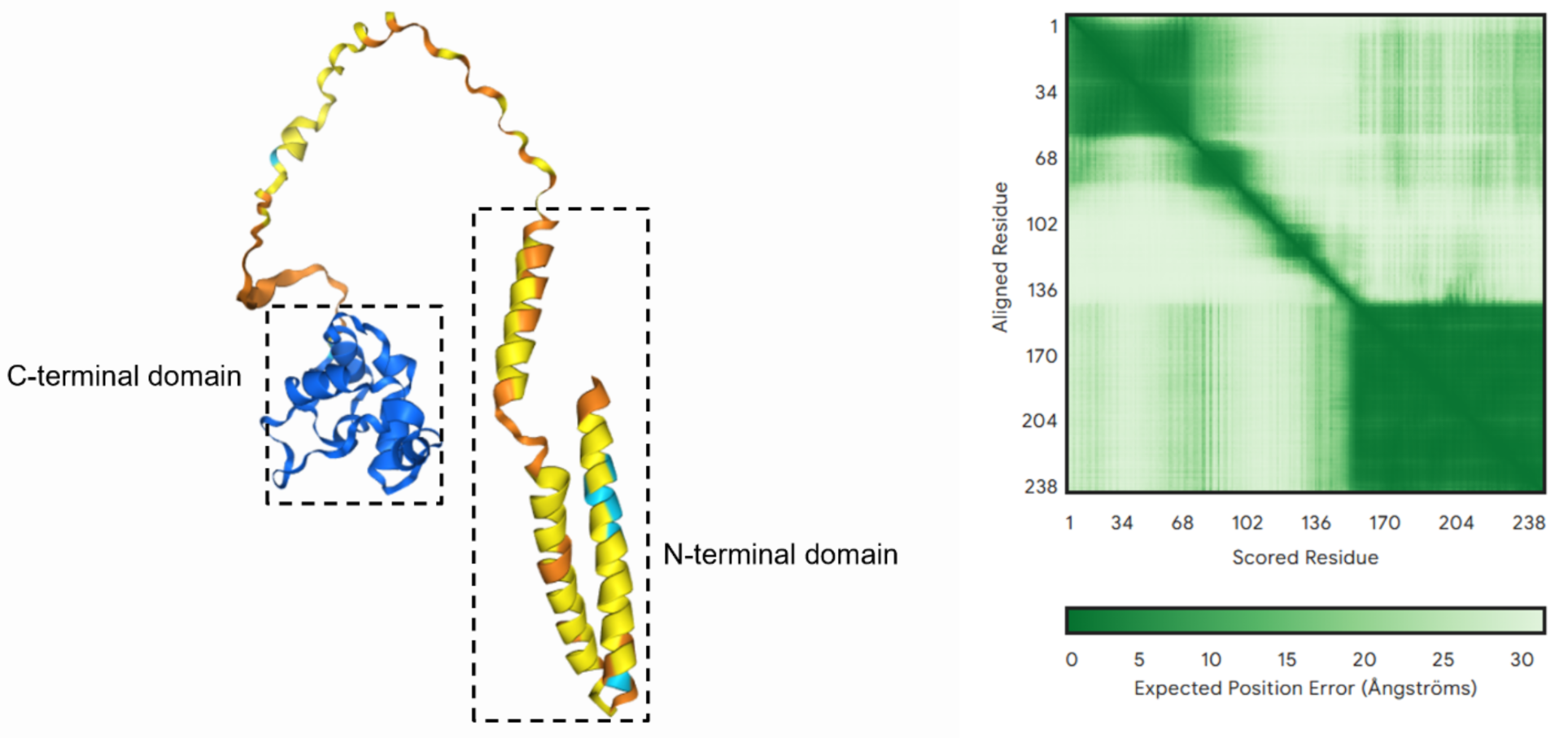
Overall structure of the full-length protein LRC predicted by AlphaFold2.

**Fig. S5.**
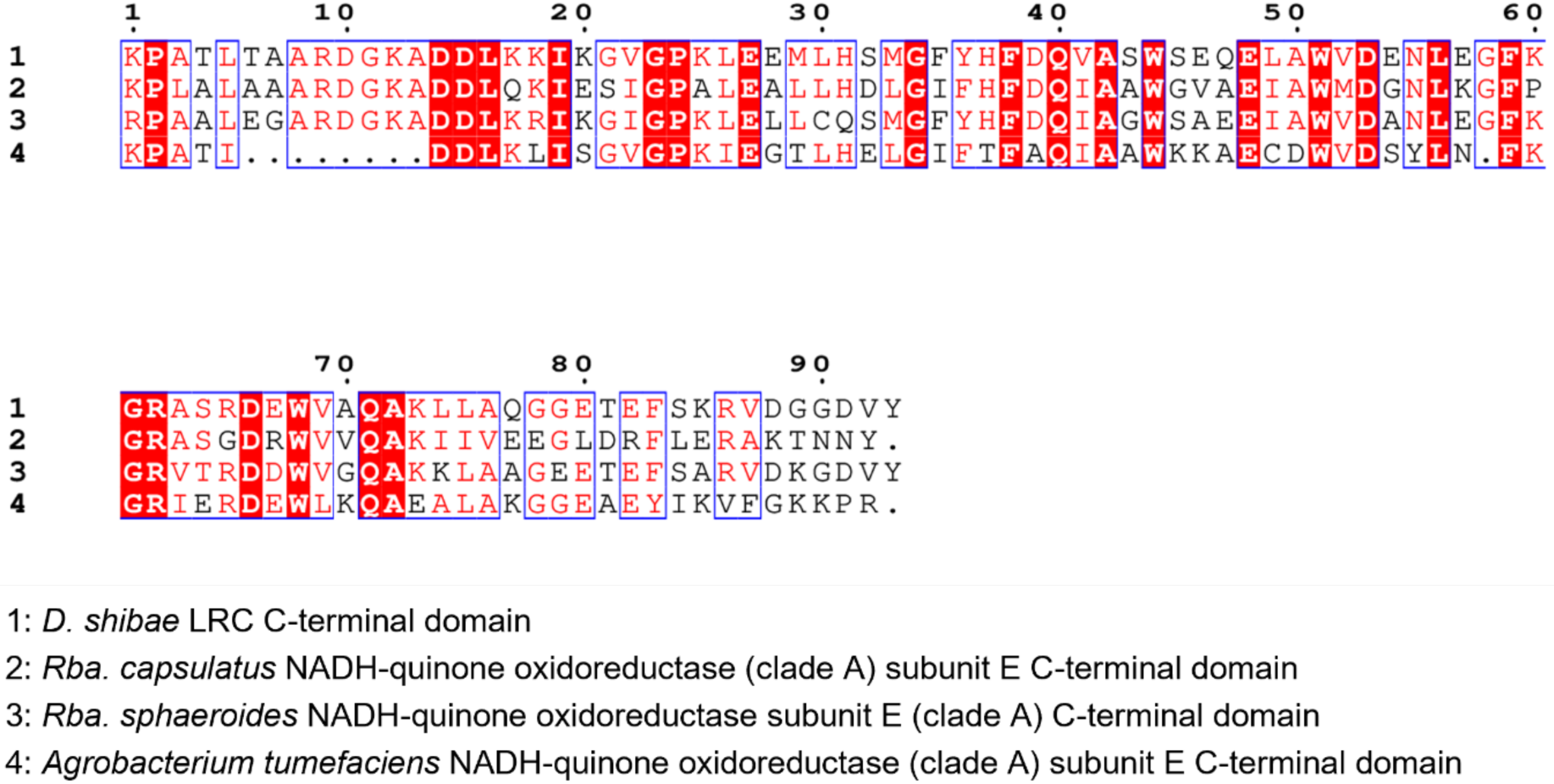
Sequence alignment of the C-terminal domain of the LRC protein from *D. shibae* and the C-terminal domains of the NADH-quinone oxidoreductase subunit E from *Rba. capsulatus*, *Rba. sphaeroides,* and *Agrobacterium tumefaciens*.

**Fig. S6.**
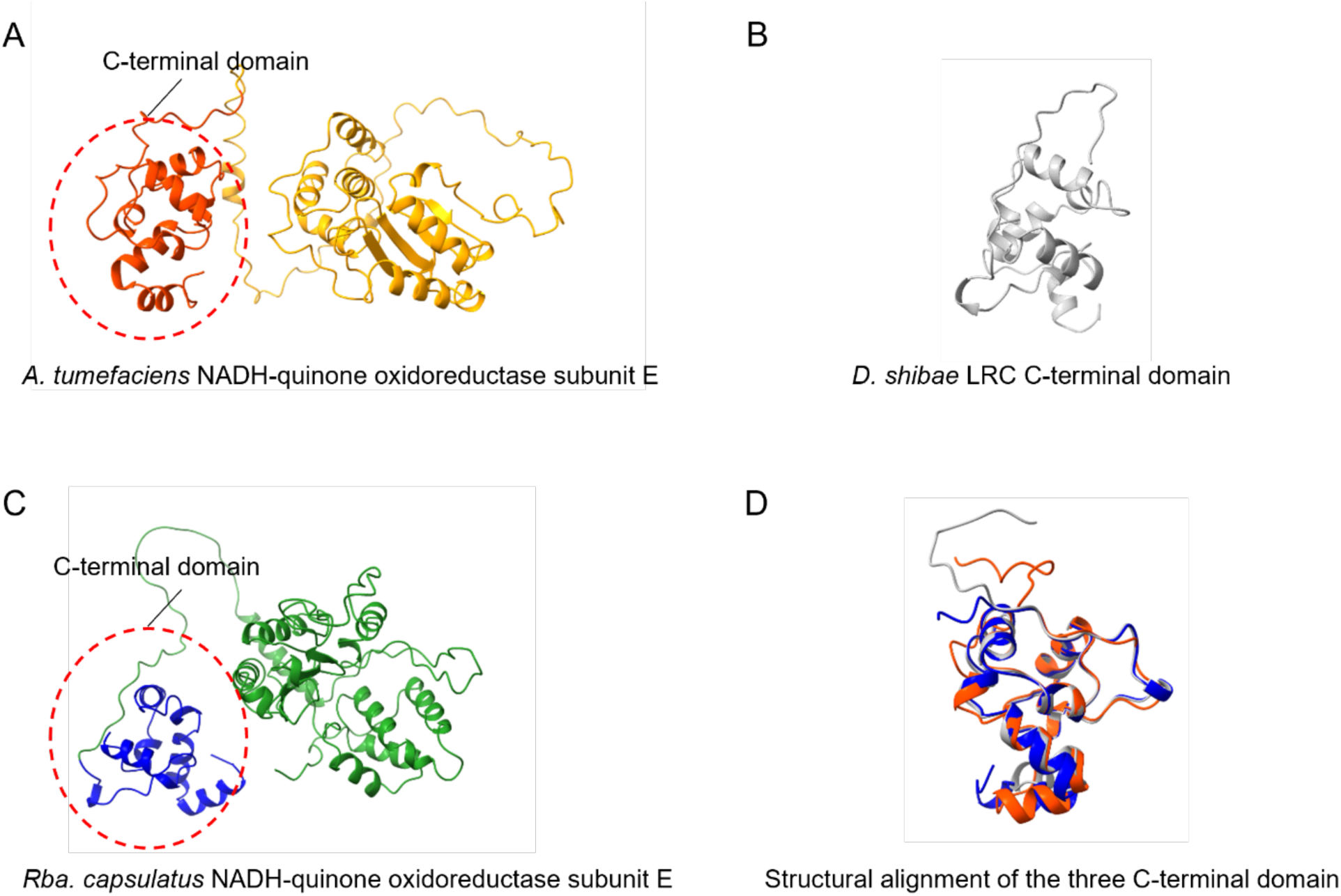
Structural comparison of the C-terminal domains of clade A NADH-quinone oxidoreductase subunit E from different bacteria and the LRC protein. (A) Structure of clade A NADH-quinone oxidoreductase subunit E from *A. tumefaciens*. (B) Structure of the LRC C-terminal domain from *D. shibae*. (C) Structure of clade A NADH-quinone oxidoreductase subunit E from *Rba. capsulatus*. (D) Structural alignment of the C-terminal domains of Clade A NADH-quinone oxidoreductase subunit E from different bacteria and the protein LRC.

**Fig. S7.**
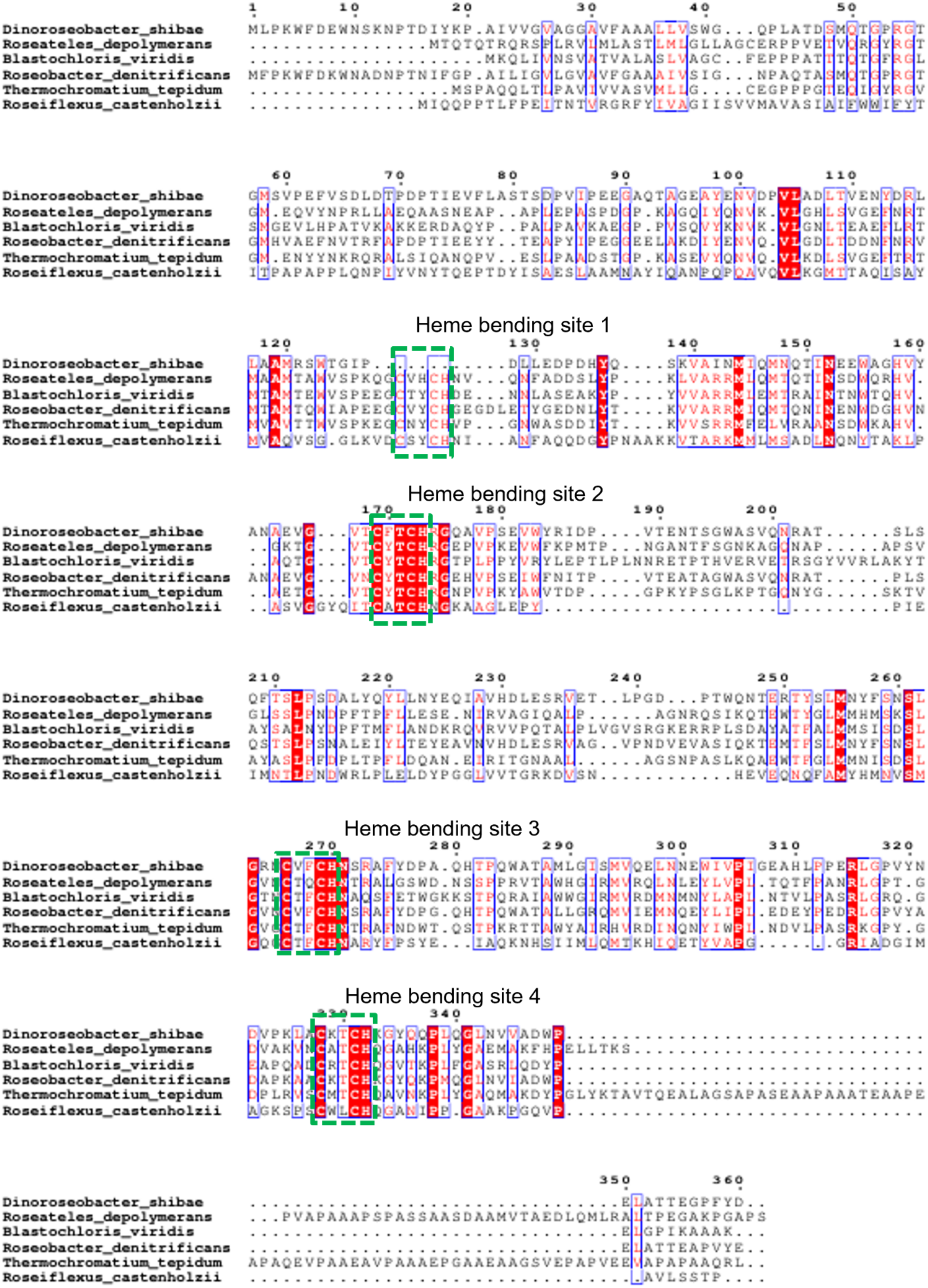
Multiple sequence alignment of cytochrome c subunits from *D. shibae* and other representative photosynthetic bacteria.

**Fig. S8.**
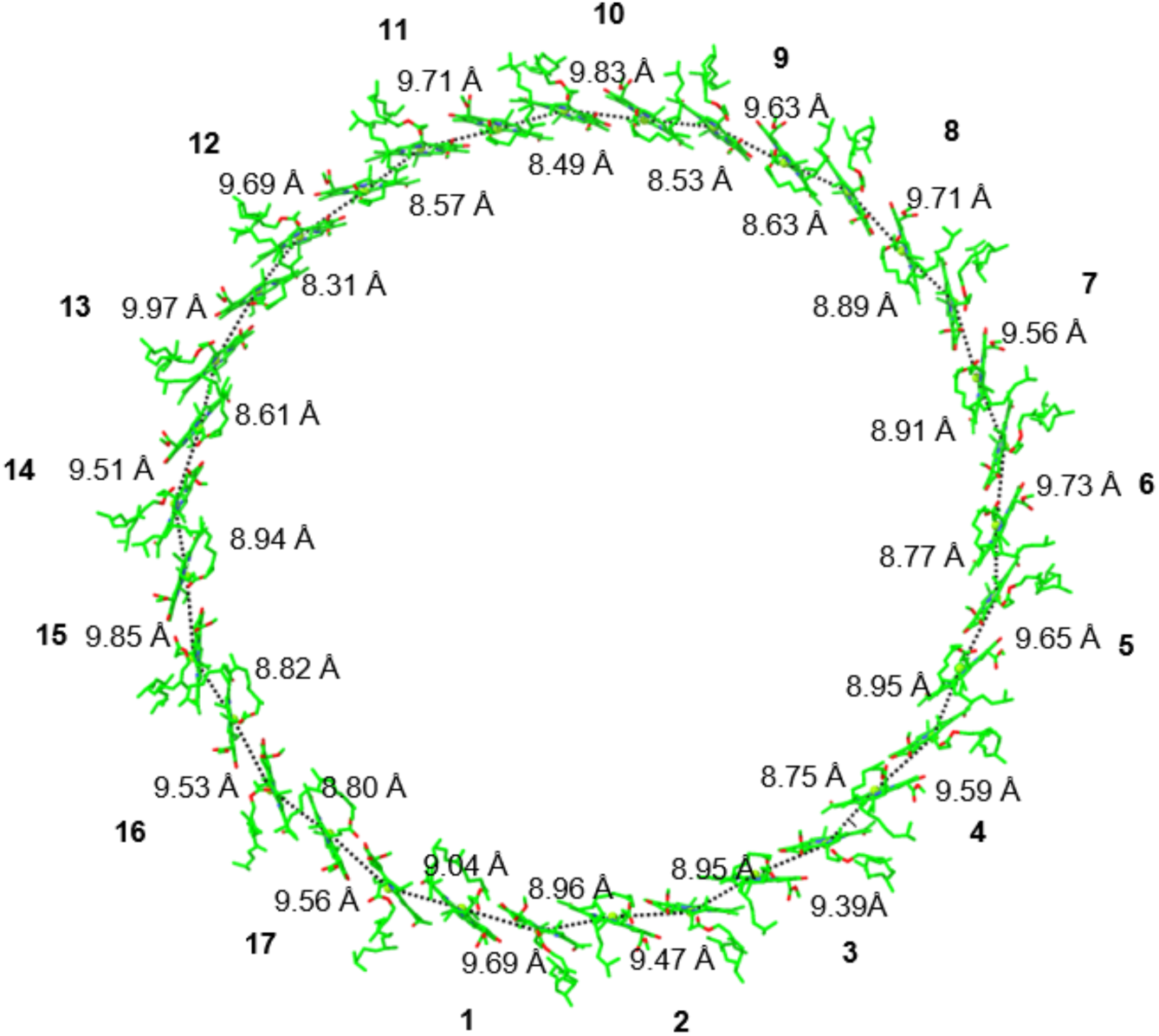
Intra-and inter-subunit Mg-Mg distances between the BChls within LH1 ring of the *D. shibae* RC–LH1 supercomplex.

**Fig. S9.**
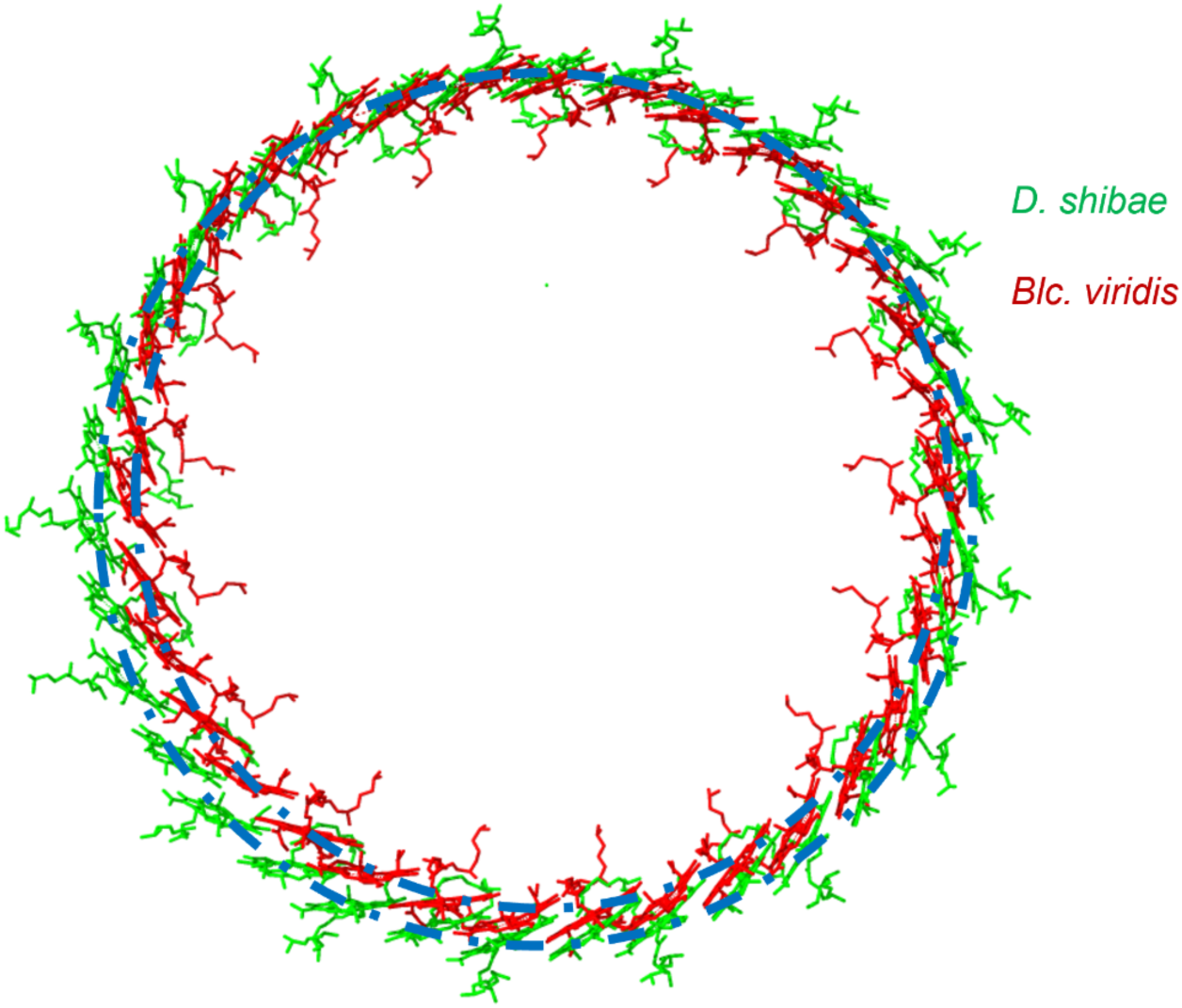
Comparison of the BChl a ring within LH1 between *D. shibae* and *Blc. viridis*.

**Fig. S10.**
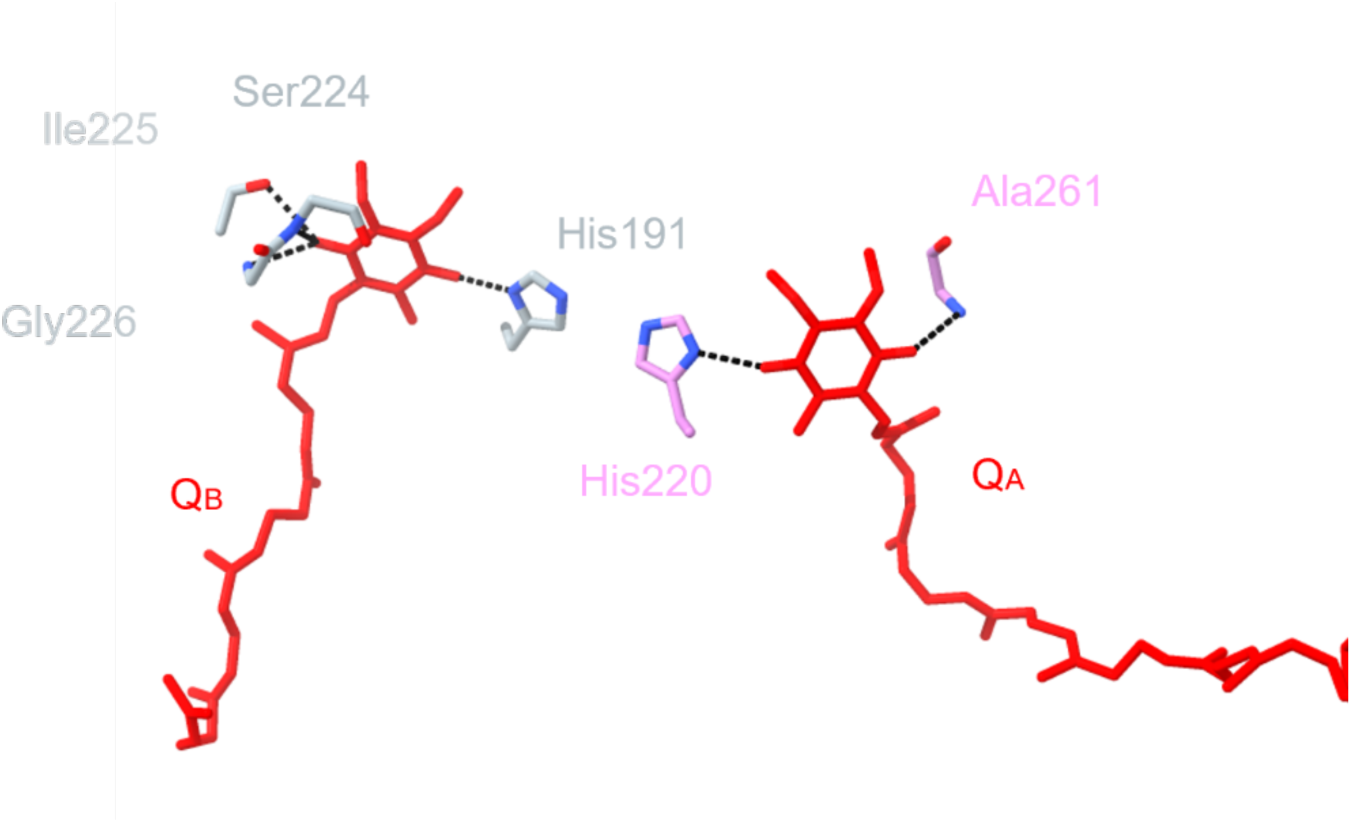
Hydrogen bond interactions involved in Q_A_ and Q_B_ binding.

**Fig. S11.**
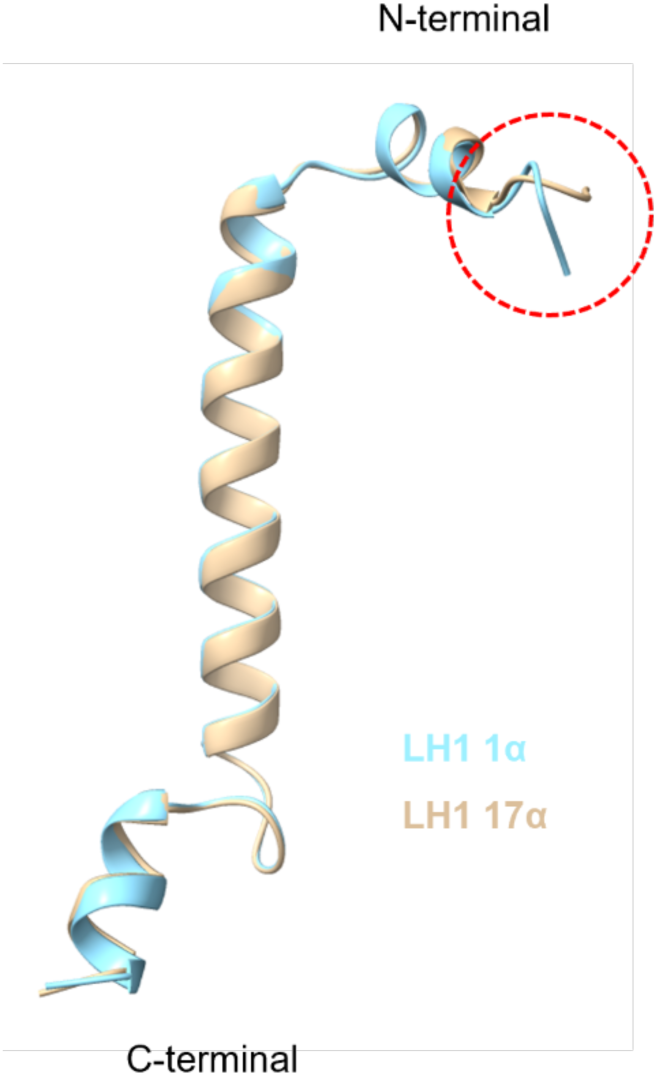
Structural alignment of LH1-1α and LH1-17α.

**Fig. S12.**
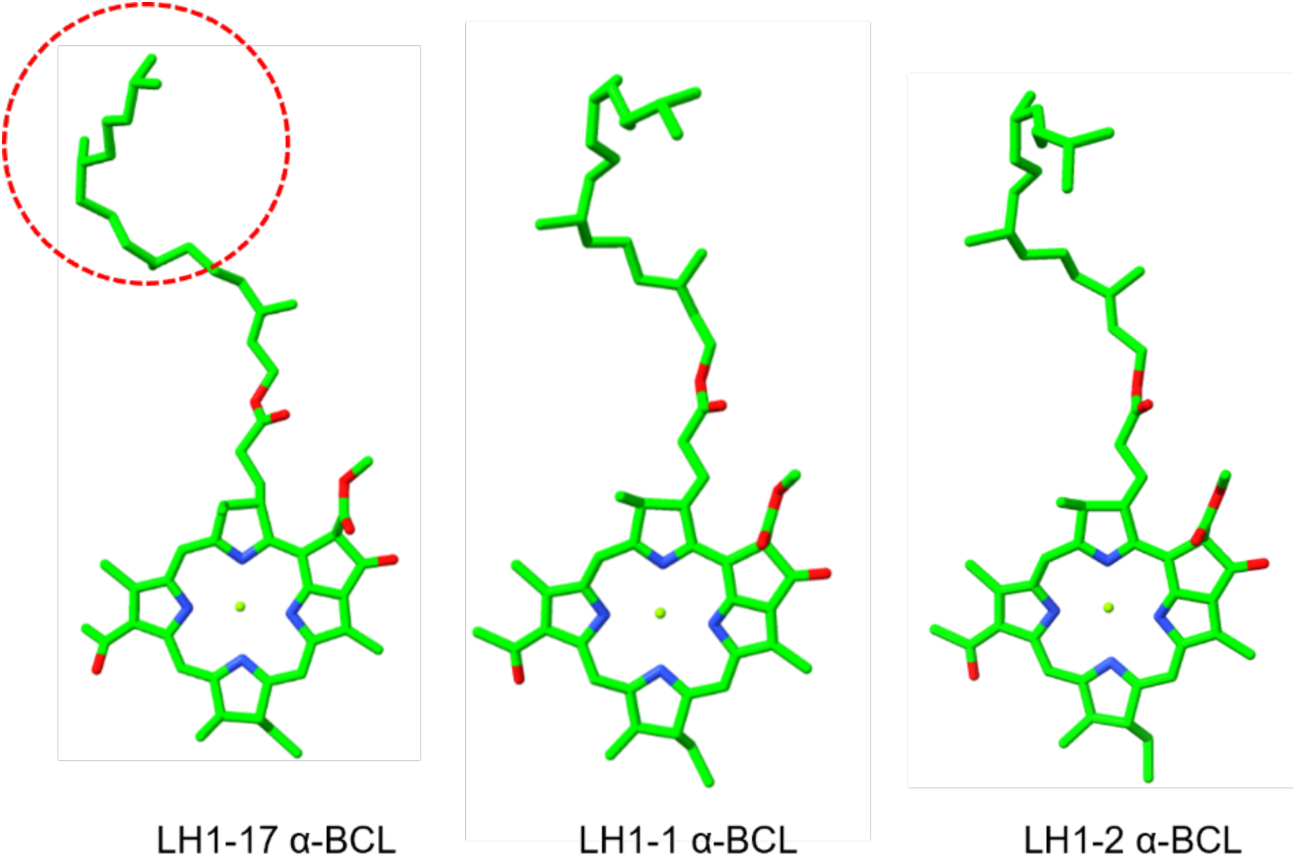
Structural comparison of BChls from different LH1 α subunits.

**Fig. S13.**
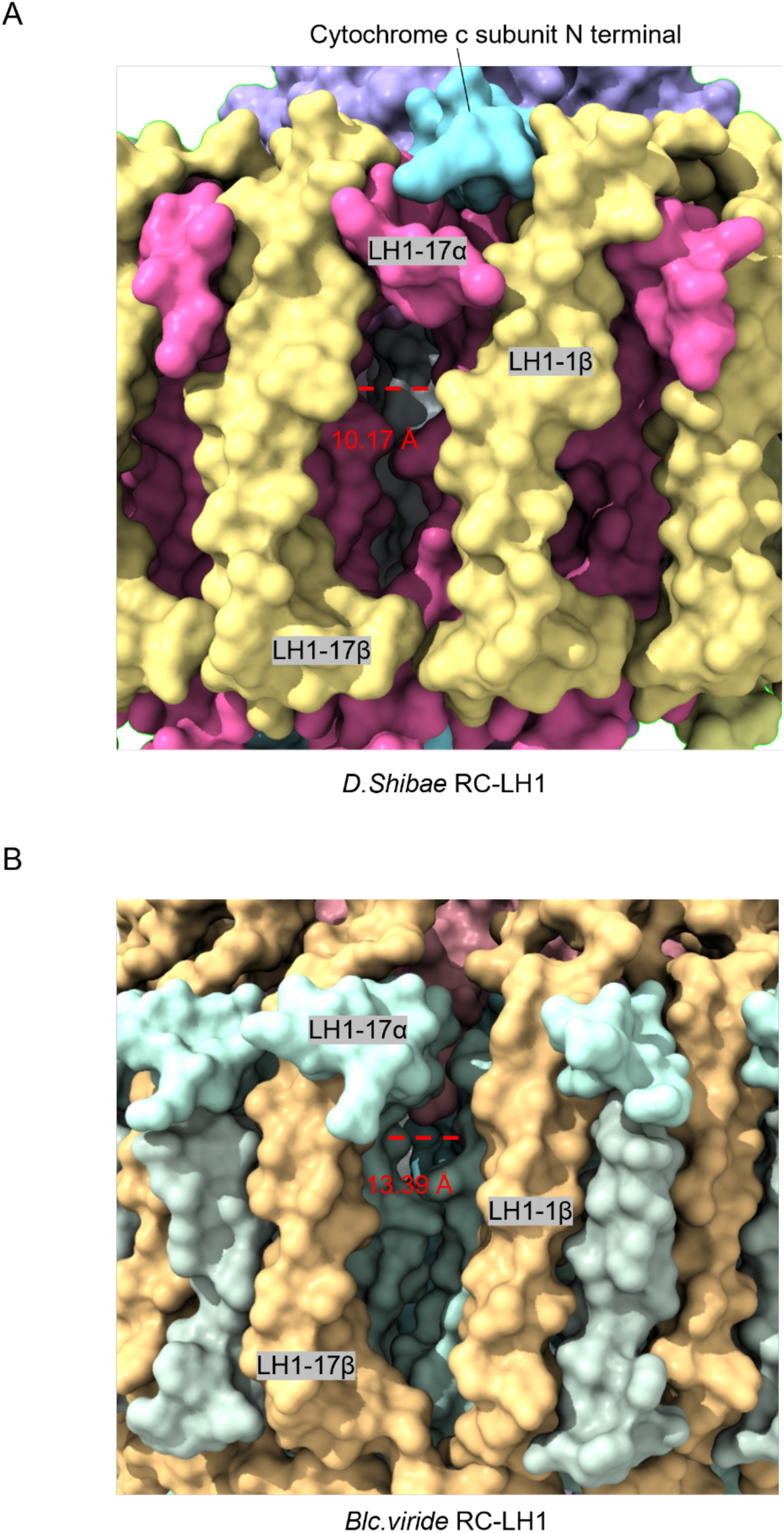
Channels for quinone and quinol diffusion across the LH1 ring in *D. shibae* RC–LH1 (A) and *Blc. viridis* RC–LH1 (B).

**Table S1.** Cryo-EM data collection, refinement and validation statistics of *D. shibae* RC–LH1 supercomplex.

|  | *D.shibae* RC-LH1<br>(EMDB-63379; PDB:<br>9LTS ) | *D.shibae* RC-LH1-A<br>(EMDB-63381; PDB:<br>9LTU ) | *D.shibae* RC-LH1-B<br>(EMDB-63382; PDB:<br>9LTV ) |
| --- | --- | --- | --- |
| **Data collection and processing** |  |  |  |
| Magnification (nominal) | 130,000 x | 130,000 x | 130,000 x |
| Voltage (kV) | 300 | 300 | 300 |
| Detector | Gatan K3 | Gatan K3 | Gatan K3 |
| Electron exposure (e<sup>-</sup>/Å<sup>2</sup>) | 40 | 40 | 40 |
| Defocus range (μm) | 0.8-1.8 | 0.8-1.8 | 0.8-1.8 |
| Pixel size (Å) | 0.97 | 0.97 | 0.97 |
| Symmetry imposed | C1 | C1 | C1 |
| Initial particle images (no.) | 1,011,021 | 1,011,021 | 1,011,021 |
| Final particle images (no.) | 123390 | 44589 | 82830 |
| FSC threshold | 0.143 | 0.143 | 0.143 |
| Map resolution (Å) | 2.49 | 2.44 | 2.75 |
| **Refinement** |  |  |  |
| Initial model used (PDB code) | 6ET5 | 6ET5 | 6ET5 |
| Model resolution (Å) | 2.87 | 2.87 | 2.87 |
| FSC threshold | 0.143 | 0.143 | 0.143 |
| **Model composition** |  |  |  |
| Non-hydrogen atoms | 28464 | 20307 | 24858 |
| Protein residues | 2901 | 2160 | 2616 |
| **Bfactor (Å<sup>2</sup>)** |  |  |  |
| Protein | 53.83 | 64.40 | 46.33 |
| Ligand | 49.93 | 63.37 | 53.99 |
| **R.m.s deviations** |  |  |  |
| Bond lengths (Å) | 0.011 | 0.44 | 0.35 |
| Bond angles (°) | 1.399 | 0.55 | 0.46 |
| **Validation** |  |  |  |
| MolProbity score | 1.41 | 1.62 | 1.44 |
| Poor rotamers (%) | 0.25 | 0.72 | 0.18 |
| **Ramachandran plot** |  |  |  |
| Favored (%) | 98.2 | 97.4 | 98.2 |
| Allowed (%) | 1.8 | 2.6 | 1.8 |
| Disallowed (%) | 0 | 0 | 0 |

**Table S2.** Hydrogen bond interactions between LRC and adjacent subunits.

| Atom1 | Atom2 | distance |
| --- | --- | --- |
| /A GLN 38 NE2 | /L TRP 266 O | 3.197 |
| /A GLN 38 NE2 | /L GLU 269 O | 3.482 |
| /u ARG 14 NH1 | /A ASP 62 O | 2.934 |
| /u ARG 14 NH2 | /A ASP 62 O | 3.205 |
| /u ARG 15 NE | /A VAL 58 O | 2.978 |
| /w ARG 15 NH1 | /A GLY 60 O | 3.190 |
| /A ARG 79 NH2 | /2 PHE 8 O | 2.709 |
| /A ARG 68 NH2 | /w ASP 12 OD2 | 3.627 |

**Table S3.**
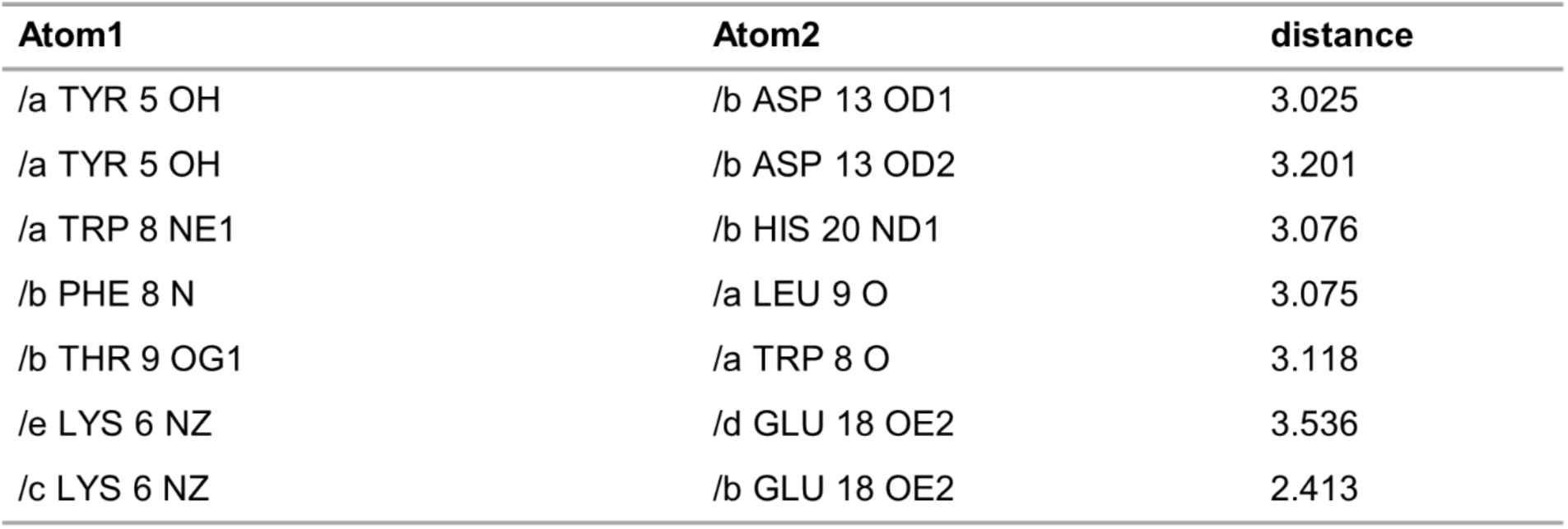
Hydrogen bond interactions within the LH1 αβ-subunit.

| Atom1 | Atom2 | distance |
| --- | --- | --- |
| /a TYR 5 OH | /b ASP 13 OD1 | 3.025 |
| /a TYR 5 OH | /b ASP 13 OD2 | 3.201 |
| /a TRP 8 NE1 | /b HIS 20 ND1 | 3.076 |
| /b PHE 8 N | /a LEU 9 O | 3.075 |
| /b THR 9 OG1 | /a TRP 8 O | 3.118 |
| /e LYS 6 NZ | /d GLU 18 OE2 | 3.536 |
| /c LYS 6 NZ | /b GLU 18 OE2 | 2.413 |

**Table S4.**
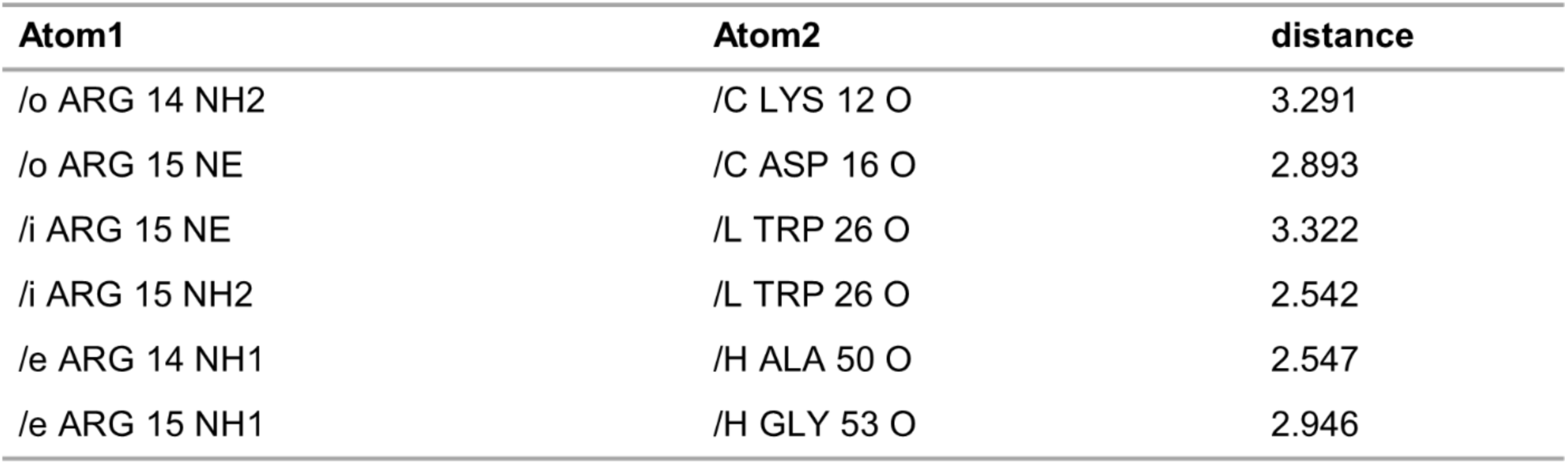
Hydrogen bond interactions between the RC and LH1 ring.

**Table S5.** Hydrogen bond interactions involved in Q_A_ and Q_B_ binding.

| Atom1 | Atom2 | distance |
| --- | --- | --- |
| /L HIS 191 ND1 | /L U10 1021 O2 | 2.340 |
| /L SER 224 OG | /L U10 1021 O5 | 2.901 |
| /L ILE 225 N | /L U10 1021 O5 | 3.156 |
| /L GLY 226 N | /L U10 1021 O5 | 3.534 |
| /M HIS 220 ND1 | /L U10 1020 O2 | 2.905 |
| /M ALA 261 N | /L U10 1020 O5 | 2.933 |

